# Proprioceptive accuracy in immersive virtual reality: A developmental perspective

**DOI:** 10.1101/760553

**Authors:** Irene Valori, Phoebe E. McKenna-Plumley, Rena Bayramova, Claudio Zandonella Callegher, Gianmarco Altoè, Teresa Farroni

## Abstract

Proprioceptive development relies on a variety of sensory inputs, among which vision is hugely dominant. Focusing on the developmental trajectory underpinning the integration of vision and proprioception, the present research explores how this integration is involved in interactions with Immersive Virtual Reality (IVR) by examining how proprioceptive accuracy is affected by *age, perception*, and *environment*. Individuals from 4 to 43 years old completed a self-turning task which asked them to manually return to a previous location with different sensory modalities available in both IVR and reality. Results were interpreted from an exploratory perspective using Bayesian model comparison analysis, which allows the phenomena to be described using probabilistic statements rather than simplified reject/not-reject decisions. The most plausible model showed that 4–8-year-old children can generally be expected to make more proprioceptive errors than older children and adults. Across age groups, proprioceptive accuracy is higher when vision is available, and is disrupted in the visual environment provided by the IVR headset. We can conclude that proprioceptive accuracy mostly develops during the first eight years of life and that it relies largely on vision. Moreover, our findings indicate that this proprioceptive accuracy can be disrupted by the use of an IVR headset.

## Introduction

From the intrauterine life, our physical, psychological, and social development progresses thanks to the interaction between our genetic profile and the environment. Sensory information from the both external world (*exteroception*) and the self (*interoception*) is detected by our emerging sensory functions. We talk about exteroception when the sensory information comes from the environment around us (e.g. sight, hearing, touch), while interoception is the perception of our body and includes “temperature, pain, itch, tickle, sensual touch, muscular and visceral sensations, vasomotor flush, hunger, thirst” (p. 655 [1]). This information, which comes from different complementary sensory modalities, has to be integrated so that we can interact with and learn from the environment. The multisensory integration that follows takes time to develop and emerges in a heterochronous pattern: we rely on the various sensory modalities to different degrees at different points in the human developmental trajectory, during which the sensory modalities interact in different ways [2]. In general, our sensory development is driven by crossmodal calibration: one accurate sensory modality can improve performance based on information delivered by another, less accurate, sensory modality [3–5].

### Proprioception: an emergent perception arising from a multisensory process

Both exteroception and interoception drive our discovery of the external world and the self. One important physical dimension of the concept of self is *proprioception*, whose definition is particularly complex and debated in the extant literature. Proprioception belongs to the somatosensory system [6] and has traditionally been defined as the “awareness of the spatial and mechanical status of the musculoskeletal framework” which includes the senses of position, movement, and balance (p. 667 [7]). From this perspective, proprioception is the awareness of the position and movement of our body in space and results from the processing of information from muscle and skin receptors. It arises from static (position) and dynamic (movement) information, and is crucial to the production of coordinated movements [8]. In general, researchers are now bypassing the concept of different unimodal sensory processing to conceive of perception as essentially multimodal, leading to multisensory interpretations of proprioception. In blind conditions, humans rely on somatosensory information to achieve proprioception, although proprioception can also emerge from vision alone. This is evidenced by studies of mirror therapy for phantom limb pain [9] that demonstrate that vision can induce proprioceptive sensations, perception of movement, touch, and body ownership, even when somatosensory input is completely absent. Similarly, as demonstrated by the rubber-hand illusion [10], visual-proprioceptive information calibrates somatosensory-proprioceptive information to create proprioception. This is why we can perceive an illusionist proprioception (of our perceived hand position) going beyond our somatosensory-proprioceptive input (of our actual hand position). Synchronous multisensory stimulation creates proprioception, while vision alone is not sufficient to induce the rubber-hand illusion when the visual input is asynchronous to the somatosensory information arising from the real hand.

These studies highlight that proprioception is not a sense like vision or touch. Rather, proprioception is a complex body consciousness which flexibly emerges from different interdependent sensory inputs, modalities, and receptors. Proprioceptive information is combined with information from the vestibular system, which detects movement of the head in space, and the visual system to give us a sense of motion and allow us to make estimates about our movements [11]. As such, it plays a vital role in everyday tasks such as self-motion.

As regards the development of proprioception, children up to two years of age tend to make significant proprioceptive errors [12]. While several studies have shown that proprioceptive competence is stably developed by eight years of age [13, 14], others support the finding of a longer developmental trajectory for proprioception, observing that 8-to 10-year-old children are less accurate than 16- to 18-year-old adolescents when making proprioceptively guided movements [15]. Moreover, some studies find improvements in proprioceptive accuracy continuing up to 24 years of age [16].

This proprioceptive development seems to be strictly dependent on visuo-proprioceptive calibration. In general, sensory organization is qualitatively different across development and across different tasks. In infancy and early childhood, vision appears dominant over somatosensory and vestibular information [17]. Between five and seven years of age, visual influence on proprioception shows non-linear developmental differences [18], although this has not yet been widely studied in a broader age ranges [17]. The developmental trajectory of proprioception may be affected by the fact that across childhood, the sections of the body change in terms of size, shape, relative location, and dynamic. Indeed, the early importance of vision over somatosensory information could be a result of the lack of reliability of somatosensory input, which is highly unstable during these childhood physical changes [2].

### IVR as a method of studying proprioception

The degree to which vision influences proprioception at different ages is an intriguing topic which can be effectively investigated in the emerging field of Immersive Virtual Reality (IVR). This tool manipulates vision and makes the user actively interact with the virtual environment, requiring actions based on proprioception. Through IVR, we can manipulate individual sources of sensory information, be they visual, vestibular, or proprioceptive, which are physiologically bound together. This makes it possible to study the contribution of these individual sensory inputs and of multisensory integration to self-perception and motor control [19]. Furthermore, it allows us to see how these individual senses contribute to proprioceptive accuracy at different developmental stages.

In IVR, “the simultaneous experience of both virtual environment and real environment often leads to new or confounded perceptual experiences” (p.71 [20]). For example, users can see themselves standing in the empty space between two mountains but, instead of falling, perceive the floor under their feet. Even with a virtual body representation (e.g. visual perception of an avatar) or without the possibility to see one’s own body, IVR can alter a user’s body schema [21]. The available literature provides some examples of how IVR affects the user’s motor activity, which relies largely on proprioception. IVR users are found to decrease their speed and take smaller steps [22] and experience greater difficulties orienting themselves in a virtual environment (VE) [23]. To orient and move in space in different environments and tasks, people can switch between reference frames related to the body (proprioception) or to the external world (e.g. vision). It has been suggested that IVR provides unexpected incongruent stimuli and induces a sensory conflict between vision and proprioception which differently affects users (e.g. sometimes causing motion sickness) depending on their dominant reliance on one of these two reference frames [24].

One of the central ways to investigate visuo-proprioceptive integration in IVR is through the study of self-motion [25]. In the area of simulated self-motion, Riecke and colleagues [26] have shown that IVR disrupts adults’ ability to perform simulated upright rotations and their judgements of these rotations. Participants’ accuracy in these rotations was markedly impaired when wearing a head-mounted display (HMD) showing them an immersive virtual environment (IVE), compared to a curved or flat screen. Despite their inaccuracy, participants subjectively rated the task as rather easy. It appears that the use of IVR or of HMDs specifically may affect proprioceptive accuracy beneath the level of awareness of the user.

Moving beyond passive self-motion, a further body of studies have examined active movements. Active self-motion involves IVEs where free movements are possible: the IVR scene changes consistently with the user’s active movement. For example, the user sees themselves walking in the virtual environment while they are physically walking in reality [27]. The ability to make active movements during the interaction with an IVR environment, even without visual landmarks, improves the perception of self-motion [28, 29]. However, despite the importance of the body senses, the physical feedback (derived, for example, from the possibility to actively walk during the virtual immersion) is not sufficient to eliminate errors in self-motion and spatial orientation while wearing an HMD [27]. These findings, taken together, show that IVR, and HMD-delivered IVR in particular, can disrupt proprioception in adults.

The studies described above primarily tested adult populations. However, there is a lack of research regarding how IVR affects proprioception, visuo-proprioceptive integration, and self-motion during development. A recent experimental study with children (8–12 years old) and adolescents (15–18 years old) provides evidence about children’s use of vision and proprioception during self-motion in IVR [30]. The authors intentionally created a mismatch between visual (visual flow) and proprioceptive feedback (active motion) during two different motor tasks: walking and throwing. They measured children’s ability to *recalibrate* (to adapt their motor actions to the provided abnormal visual input) and *re-adapt* to the normal characteristics of the real environment. As with adults in previous studies [31, 32], children and adolescents showed the ability to recalibrate in a few minutes. The authors found just one age-related difference, in regard to the rate of re-adaptation. In the throwing task, children re-adapted to reality significantly more slowly than adolescents, demonstrating more pronounced post-exposure effects. The mismatch between visual and proprioceptive information appeared to have a more enduring effect on children. Although this finding must be interpreted with caution, it could be a first indication of age-related differences in motor learning in IVR. These findings indicate that the motor performance of children, more so than adolescents, could be modified by interaction with IVR environments. This could have meaningful implications for fields such as IVR rehabilitation, therapy, and education, suggesting that IVR interventions could be more effective early in life.

With concern to multisensory integration, a recent study used IVR to decouple visual information from self-motion and investigate whether adults and 10- and year-old children can optimally integrate visual and self-motion proprioceptive cues [33]. A HMD was used to make participants learn a two-legged path either in darkness (“only proprioception”), in a virtual room (“vision + proprioception”), or staying stationary while viewing a pre-recorded video of walking the path in the virtual room (“only vision”). Participants then reproduced this path in darkness. In contrast to what was expected, the authors found that adults failed to optimally integrate visual and proprioceptive cues to improve path reproduction. However, children did integrate these cues to improve their performance. This study demonstrates that HMD training that includes vision and proprioception can be effective at calibrating self-motion for children even if it is not for adults. The authors suggest that this may be because children cannot help but rely on visual cues in spatial tasks even when the nature of the task does not require it. The authors do not explain the results with respect to the use of IVR, or specifically by considering IVR as a tool which requires a particular form of sensory processing. We previously discussed findings demonstrating that HMDs disrupt proprioception, which adults and children rely on in different ways. It may be the case that IVR imparts different effects on adults’ and children’s performance. We could speculate that, if IVR causes some sort of conflict between vision and proprioception, adults’ lack of multisensory integration in these environments could be due to their reliance on proprioception and ability to ignore visual cues. Visual cues would be perceived as irrelevant for motor tasks, because they would be in conflict with proprioceptive information. Since this ability to ignore irrelevant visual cues seems not to be mature in children [34], they could benefit from IVR motor training because they would still be using vision to calibrate their less accurate proprioception. It is only recently that the field of IVR research is beginning to focus on the developing child to study developmental differences in relation to their interaction with IVR environments. Thus far, IVR technology has been primarily used with children for educational, pain distraction, and assessment purposes [35]. Further research is needed to investigate how the sensory-motor interaction with an IVR environment changes depending on age. Given that children and adults differ in their sensory-motor functioning, research should investigate how IVR interacts with and affects the childhood developmental trajectory with respect to use of vision, proprioception, and other sensory cues in the ability to accurately execute self-motion.

### Statistical approach for exploratory investigations: Bayesian model comparison

Given the lack of evidence concerning the complex interaction between developmental stages, visuo-proprioceptive integration, and IVR environments, exploratory studies are needed and can benefit from assuming a model comparison approach. Model comparison allows for the selection of the most plausible model given data and a set of candidate models [36]. Firstly, the different research hypotheses are formalized as statistical models. Subsequently, the obtained models are compared in terms of statistical evidence (i.e. support by the obtained data), using information criteria [37]. Information criteria enables the evaluation of models considering the trade-off between parsimony and goodness-of-fit [38]: as the complexity of the model increases (i.e. more parameters), the fit to the data increases as well, but generalizability (i.e. ability to predict new data) decreases. The researchers’ aim is to find the right balance between fit and generalizability in order to describe, with a statistical model, the important features of the studied phenomenon, but not the random noise of the observed data.

A Bayesian approach is a valid alternative to the traditional frequentist approach [39, 40], allowing researchers to accurately estimate complex models that otherwise would fail to converge (i.e. unreliable results) in a traditional frequentist approach [41, 42]. Without going into philosophical reasons, which are beyond the scope of the present paper (if interested, consider [43]), Bayesian inference has some unique elements that make the meaning and interpretation of the results different from the classical frequentist approach [44]. In particular, in the Bayesian approach, parameters are estimated using probability distributions (i.e. a range of possible values) and not a single point estimate (i.e. a single value). Bayesian inference has three main ingredients [45]: (1) *Priors*, the probability distributions of possible parameter values considering the information available before conducting the experiment; (2) *Likelihood*, the information given by the observed data about the probability distributions of possible parameter values; (3) *Posteriors*, the resulting probability distributions of possible parameter values, obtained by combining Priors and Likelihood through Bayes’ Theorem. As a result, a Bayesian approach assesses the variability (i.e. uncertainty) of parameter estimates and provides associated inferences via 95% Bayesian Credible Intervals (BCIs), the range of most credible parameter values given the prior distribution and the observed data. Thus, a Bayesian approach allows researchers to describe the phenomenon of interest through probabilistic statements, rather than a series of simplified reject/do-not-reject dichotomous decisions typically used in the null hypothesis significance testing approach [36].

### Research goals and hypotheses

The aim of the present study is to investigate the extent to which the reliability of visual information aids proprioceptive-based self-motion accuracy across the human developmental trajectory. We also aim to explore whether HMD-delivered IVR environments, compared to equivalent real environments, affect proprioceptive accuracy. Given that findings in the area of multisensory interaction with IVR across development are still conflicting and unexplained with respect to the use of HMDs, the current study seeks to clarify how using an HMD affects children’s and adults’ self-motion performance, and how these effects could be related to the reliability of the provided visual and proprioceptive information. Research has broadly considered the computer side of IVR features affecting human-computer interaction, but there is a lack of research investigating how individual characteristics of users interact with IVR environments. To compare performances in reality and IVR, all sensory conditions being equal, would clarify the role of both sensory manipulation and IVR. How might different users, with different levels of multisensory functioning, interact with IVR? The present study explores this question, examining how IVR differs from reality in affecting visuo-proprioceptive integration in adults and children at different developmental stages. Furthermore, the study aims to open new avenues of analysis in this area of research by using a model comparison approach to analyze each hypothesis.

Based on the extant literature described in the introductory section of this work, we hypothesized that children’s proprioceptive accuracy would be globally lower than that of adults, but that children would be less impaired than adults by the disruption of proprioception. We further hypothesized that IVR would disrupt proprioception and impact proprioceptive accuracy more in adults than children.

## Materials and Methos

### Participants

In order to capture a range of developmental stages, we included primary and secondary school-aged children and adults. We collected data from young children aged from 4 to 8 years old, and older children aged from 9 to 15 years old. This distinction was made to clarify contradictory findings about how long it takes to develop stable proprioceptive accuracy (as described in section 2.2). With regard to the adult group, we included participants within the age range of 18 to 45 years. We excluded older participants based on literature reporting deterioration of proprioceptive accuracy with advancing age. This deterioration effect has been found from middle age, with studies indicating changes beginning from the age of 40 to 60 [46, 47]. For this study, we collected data from 55 participants. In line with our a priori exclusion criteria, we excluded six subjects who reported that they had received a diagnosis for any kind of neuropsychological, sensory, or learning disorder from the final analysis. The final sample included 49 participants, distributed across age groups as follows:

- 13 young children between the ages of 4 and 8 years (*M*_*age*_ = 7.1, *SD* = 1.2 years)
- 13 older children between the ages of 9 and 15 years (*M*_*age*_ = 11.3, *SD* = 2.0 years)
- 23 adults between the ages of 20 and 43 years (*M*_*age*_ = 32.4, *SD* = 6.6 years)

In a within-subjects design, all participants were exposed to all conditions in a randomized order.

### Materials and set-up

We designed and built a testing room in which different sensory stimulations could be provided and the availability of visual and proprioceptive information could be manipulated while completely excluding unwanted external stimuli (Fig 1). In the centre of the room, we fixed a customized swivel chair on a round platform to the floor. The round platform did not provide any proprioceptive or visual cues about the degree of rotation the participant made on the chair (Fig 2A). A 360° protractor under the seat was visible via a dedicated camera which allowed the measurement of the degree of each rotation. One 50 cm white LED strip (12V DC, 24 Watt per meter) allowed sufficient illumination for a clear and realistic visual experience of the room. One UV lamp (E27 26W) was used to obscure other visual stimuli such that the white clouds on the walls were the only visual cues available. With the UV light on, participants were asked to wear a black poncho which covered their bodies, making them not visible (Fig 2B). One infrared LED spotlight (BIG BARGAIN BW103) enabled clear video recordings of the inside of the room even when it was completely in darkness. This light system was anchored to the ceiling, over participants’ heads, and was covered by a black panel which prevented participants from directly seeing the lights.

**Fig 1.**
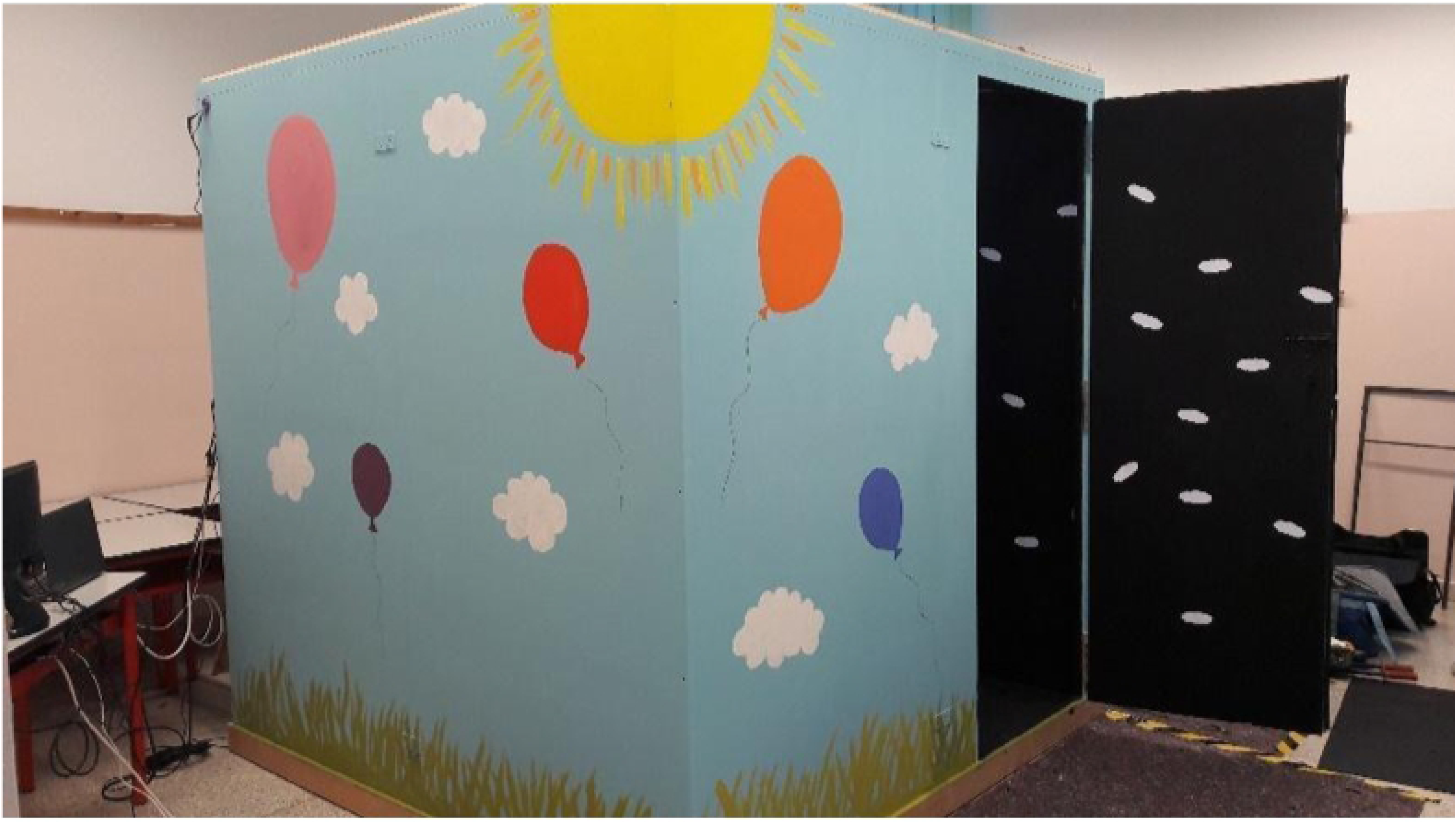
Experimental room. The room measured 2 × 2 meters and was soundproof, with black interior walls and equal numbers of white clouds randomly fixed on each wall. The external walls were painted with a child-friendly landscape which has been designed to encourage children to enter the room.

**Fig 2.**
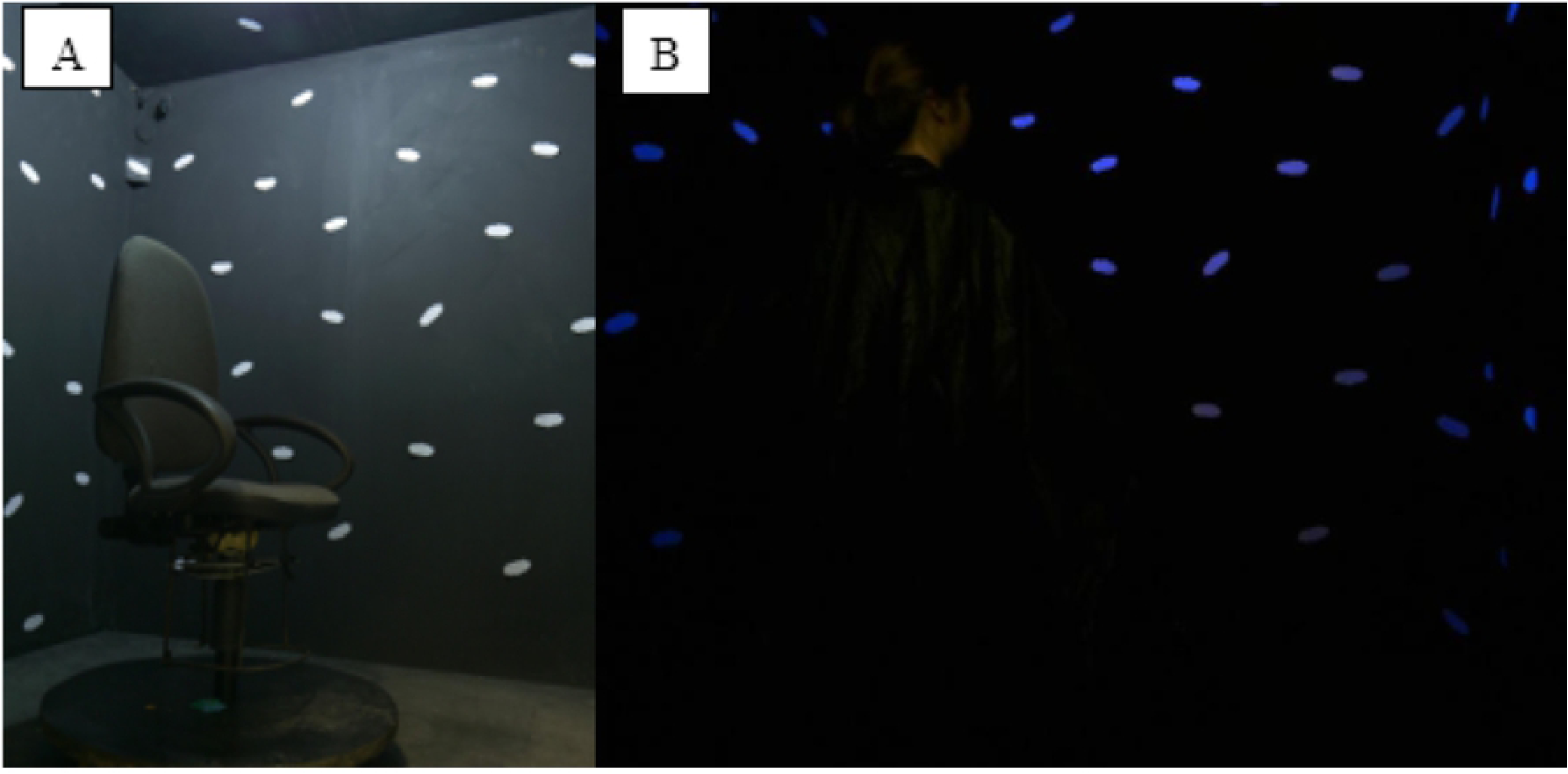
Experimental room, interior. A: The swivel chair in a visuo-proprioceptive real environment. B: A participant wearing the black poncho in a vision-only real environment (B).

We provided the IVR simulation through a VR headset (Head Mounted Display [HMD]). We used Oculus Gear VR 2016, 101° FOV, 345 g weight, interfaced with a Samsung Galaxy S7 (ANDROID 8.0.0 operating system).

A NIKON camera KeyMission 360 was used to create 360° images of the room and to build the IVR environments. The room was monitored via one USB 2.0 DirectShow webcam, and one USB 2.0 DirectShow webcam with integrated infrared LED.

To monitor the video recordings and VR simulations, we used a SATELLITE Z30-B, Windows 10, 64bit, Intel Core i5-5200U CPU @ 2.20 Ghz,8,0 GB RAM, Intel HD Graphics 5500. The communication between people inside and outside the room was enabled via a system of USB speaker, microphone, headphones, and one USB soundcard. The VR server application developed for this experiment is an Android application with VR environments, developed in Unity. A remote interface, also developed in Unity for Windows or Android OS, allowed experimenters to control the VR server application. A software for audio-video recording and real-time communication was developed in TouchDesigner.

## Procedure

Adult participants were welcomed into the lab and asked to sign a consent form. Parents of children were asked to sign the form on their child’s behalf. The study was approved by the Ethics Committee of Psychology Research, University of Padua. At least two experimenters conducted the experiment. On commencing the experiment, participants were asked to sit on the swivel chair which was fixed in the middle of the recording area inside the room. The first experimenter would close the door and stay inside near the participant for the duration of the experiment. The second experimenter managed the experiment: he/she switched the lights on and off, changed the visual stimuli which were presented through the HMD, and controlled the video recording of the experiment. He/she was outside the room, monitoring the video feed, and giving verbal instructions to the first experimenter and to the participants. The room is soundproof but the second experimenter could communicate with the people inside through a microphone. The participant and experimenter inside the room could hear the second experimenter through a system of speakers set up under the swivel chair. During the experimental task (described below in the following paragraph), the first experimenter managed the passive rotation and remained silent behind the participant, providing no visual or auditory cues. The second experimenter followed previously established verbal instructions which were consistent across participants.

### Experimental task

We adopted a self-turn paradigm in which the experimenter rotates the chair a certain degree (passive rotation) from a *start position* to an *end position*. After each passive rotation, participants were asked to rotate back to the start position (active rotation). The position at which the participant stopped their active rotation is recorded as the *return position*. All participants performed 12 trials across 6 conditions. For each condition, the passive rotation was done once to the right (clockwise) and once to the left (counterclockwise). For each condition, one passive rotation was approximately 180 degrees and the other was approximately 90 degrees. During the passive rotation, participants kept their feet on a footrest which rotated with the chair. In this way, they could not make steps while being rotated, and could not simply count the number of steps to make active rotations. To perform the active rotations, participants could use their feet on the still platform under the chair to move themselves. Some authors suggest that vestibular information is primarily involved when perceiving the amount of *passive rotation*, and proprioceptive information is primarily involved when performing an *active rotation* [48]. In our task, during the encoding phase (passive rotation), vestibular information is always available, while proprioception is not. During the recall phase (active rotation), both vestibular and proprioceptive information are available. In each experimental condition, the same vestibular and visual information can be used to both encode and recall the *start position*. Proprioception has to be used only during the recall phase, emerging from the other sensory information. Proprioception is considered as the accuracy measure in our task in line with procedures aimed at assessing proprioception in the extant literature [49–51].

### Measures of task performance

The proprioceptive accuracy of self-turn performances was calculated in terms of error as the absolute difference between the *start position* (from which the experimenter started the passive rotation) and the *return position* (in which the participant stopped the active rotation). In this way, greater values indicated a less accurate performance, where a value of 0 would indicate that the participant actively rotated back to the exact start position, and a value of 100 would indicate that the participant actively rotated back to a position that was 100 degrees away from the start position.

Proprioceptive accuracy was manually measured during an offline coding of the video recording. The video shows two matched recordings of both the entire room (with the participant and the first experimenter in frame) and the protractor positioned under the seat of the swivel chair. A vertical green line was superimposed on the protractor image to facilitate detection of the specific degree of each rotation. Two independent evaluators coded the videos and entered the start and return positions in the dataset. Values which were divergent for more than two degrees were a priori considered disagreement values. A third coder examined the video records of the disagreement values to make the final decision. In case of a disagreement value, the third coder’s value was used instead of the value that differed most from the third coder’s value. We obtained a dataset with two codings for each data. We evaluated the intercoder agreement by conducting an intra-class correlation (ICC), which is one of the most commonly used statistics for assessing inter-rater reliability (IRR) for ratio variables [52]. From the dataset which combines the two codings, we obtained a final dataset with the average of the two values. We carried out the data analysis on this final dataset.

### Conditions

The order of conditions was randomized. Participants performed blocks of two trials per condition. There were three conditions in a real environment (R) and three conditions in an immersive virtual reality (IVR) environment. In each of these two blocks, one blind condition removed all visual information such that only proprioceptive information could be used (P), one condition limited the access to visual landmarks (removing visual information about the body and corners of the room while retaining the use of vision) in order to disrupt proprioception (V), and one condition allowed the participant to access reliable visual and proprioceptive information (VP). Several studies have explored the extent to which people benefit from visual landmarks to calibrate and aid proprioceptive tasks while self-turning and it seems that different kinds of visual landmarks could be more or less useful for proprioception in different environments and tasks. In a real environment, after being disorientated by a passive rotation, people could still detect the position of global landmarks (the room’s corners), while making huge errors locating surrounding objects [53]. In a HMD-delivered virtual environment, users’ self-motion did not benefit so much from global landmarks [54]. We aimed to control whether the rotation direction and amplitude would affect performance. For this purpose, each condition was performed twice: the passive rotation was made in both directions (clockwise and counterclockwise), and with two angle amplitudes (90 and 180 degrees). As the passive rotation was manually performed by the experimenter, perfect accuracy in reaching 90 and 180 degrees was not possible. Given the variability in the actual passive rotations, we considered Amplitude as a continuous variable. We labelled the direction conditions “R” (right) for the clockwise condition and “L” (left) for the counterclockwise condition. We counterbalanced within-subjects the possible interaction effect of Direction Learning, beginning 50% of conditions with “R” and the other 50% with “L”. We labelled the amplitude conditions “A” for the 180-degree condition and “B” for the 90-degree condition. Fifty percent of each direction condition had a 180-degree amplitude and the other 50% had a 90-degree amplitude. The direction order is RLLRRLLRRLLR; and the amplitude order is ABABABABABAB. We labelled the conditions by number from one to six. As such, we had, for example, sequences labelled: 1RA-1LB-2LA-2RB, and so on. We counterbalanced the amplitude order between subjects. We tested the ABAB sequence in 50% of subjects and the BABA sequence in 50% of subjects.

The experimental conditions are as follows:

- R_P (Reality; only proprioception, no visual information available)
- R_V (Reality; only vision: low external visual landmarks with no first-person view of the body or room corners in order to disrupt proprioception).
- R_VP (Reality; vision and proprioception are available; first-person view of the body, room corners, and clouds are visually available)
- IVR_P (Immersive Virtual Reality; only proprioception, no visual information available)
- IVR_V (Immersive Virtual Reality; only vision: low external visual landmarks with no first-person view of the body or room corners in order to disrupt proprioception)
- IVR_VP (Immersive Virtual Reality; vision and proprioception are available; room corners and clouds are visually available, although first-person view of the body is not)

### Statistical approach

In order to explore how age, sensory conditions, and environmental conditions interact to affect proprioceptive accuracy, a model comparison approach was used. Firstly, each research hypothesis was formalized as a statistical model. Subsequently, the obtained models were compared in terms of statistical evidence (i.e. support by the data) using information criteria [37].

Given the complex structure of the data, Bayesian generalized mixed-effects models were used [39, 55]. Specifically, data were characterized by: (1) a continuous non-normally distributed dependent variable (i.e. rotation error); (2) a between-subject factor (i.e. Age); (3) within-subject factors (i.e. Perception condition and Environment condition); (4) a quantitative independent variable (i.e. rotation Amplitude). Mixed-effects models allow us to take into account the repeated measures design of the experiment (i.e. observations nested within subjects). Thus, participants were treated as random effects, with random intercepts that account for interpersonal variability, while the other variables are considered as fixed effects. Gamma distribution, with logarithmic link function, was specified as the family distribution of the generalized mixed-models. Gamma distribution is advised in the case of positively skewed, non-negative data, when the variances are expected to be proportional to the square of the means [56]. These conditions are respected by our dependent variable: we only have positive values, with a positive skewed distribution, and we expect a greater variability of the possible results as the model predicted mean increases (i.e. a greater dispersion of subjects’ scores when greater mean values are predicted by the model).

Analyses were conducted with the R software version 3.5.1 [57]. Models were estimated using the R package *‘brms’* [58] which is based on STAN programming language [59, 60] and employs the No-U-Turn Sampler (NUTS; [61]), an extension of Hamiltonian Monte Carlo [62]. All our models used default prior specification of the R package *‘brms’* [58]. Detailed prior specifications are reported in the supplemental online material. These priors are considered non-informative since they leave the posterior distributions to be mostly influenced by the observed data rather than by prior information. Each model was estimated using 6 independent chains of 8,000 iterations with a “warm-up” period of 2,000 iterations, resulting in 36,000 usable samples.

Convergence was evaluated via visual inspection of the trace plots (i.e. sampling chains) and R-hat diagnostic criteria [63]. All tested models showed satisfactory convergence with all R-hat ≤ 1.0008, where values close to 1 indicate convergence, and none exceeding the 1.100 proposed threshold for convergence [39]. All R-hat values and trace plots are reported in the supplemental online material.

The Watanabe-Akaike Information Criterion (WAIC; [64, 65]) was used as information criteria to select the most plausible model among the tested models, given the data. WAIC is the corresponding Bayesian version of the commonly used Akaike information criterion (AIC; [66]). WAIC-weights were computed to present the probability of each model of making the best predictions on new data, conditional on the set of models considered [36]. This allows for the comparison of models with a continuous informative measure of evidence. Finally, the most plausible model was interpreted considering the estimated posterior parameter distributions. Main effects and interaction effects were evaluated using planned comparison and graphical representations of the predicted values by model.

## Results

### Descriptives

Proprioceptive accuracy was manually measured during an offline coding of the video recording. Independent raters coded for the degree values indicating start, end, and return positions of each rotation. Based on these values, we calculated the amplitude of passive rotations and proprioceptive errors of active rotations. On these start, end, and return position values, the intra-class correlation index (ICC) has been calculated to evaluate the inter-coder reliability. The analysis estimates an ICC = .99. This nearly perfect inter-coder agreement derives from the small mean difference between the two coders’ values, within the huge range of possible values (0/360). In fact, the mean difference between coder A and coder B is minimal (Mean_*A*−*B*_ < .16).

Out of the 49 participants, 43 subjects completed the task in all 12 conditions, 4 subjects completed 11 conditions, 1 subject completed 10 conditions, and 1 subject completed 8 conditions. This failure to complete all conditions with some participants was due to technical problems which occurred with the experimental apparatus. Thus, the final data consist of 578 observations nested in 49 subjects. The number of observations in each condition is reported in Table 3 in S1 Supplemental Materials.

**Table 1.**
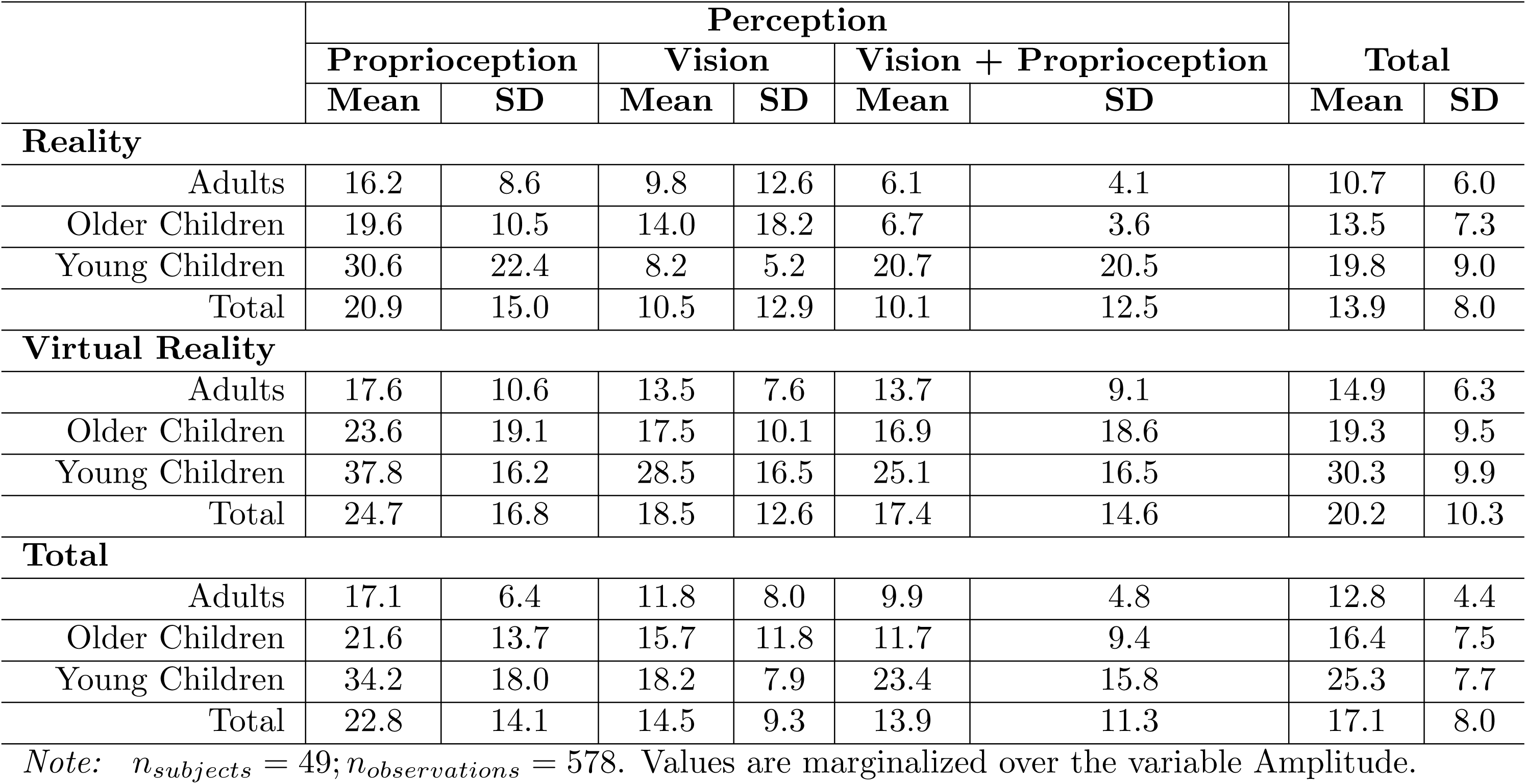
Descriptive statistics. Means and standard deviations of self-turn error according to age and the experimental conditions.

|  | Perception |  |  |  |  |  | Total |  |
| --- | --- | --- | --- | --- | --- | --- | --- | --- |
|  | Proprioception |  | Vision |  | Vision + Proprioception |  |  |  |
|  | Mean | SD | Mean | SD | Mean | SD | Mean | SD |
| Reality |  |  |  |  |  |  |  |  |
| Adults | 16.2 | 8.6 | 9.8 | 12.6 | 6.1 | 4.1 | 10.7 | 6.0 |
| Older Children | 19.6 | 10.5 | 14.0 | 18.2 | 6.7 | 3.6 | 13.5 | 7.3 |
| Young Children | 30.6 | 22.4 | 8.2 | 5.2 | 20.7 | 20.5 | 19.8 | 9.0 |
| Total | 20.9 | 15.0 | 10.5 | 12.9 | 10.1 | 12.5 | 13.9 | 8.0 |
| Virtual Reality |  |  |  |  |  |  |  |  |
| Adults | 17.6 | 10.6 | 13.5 | 7.6 | 13.7 | 9.1 | 14.9 | 6.3 |
| Older Children | 23.6 | 19.1 | 17.5 | 10.1 | 16.9 | 18.6 | 19.3 | 9.5 |
| Young Children | 37.8 | 16.2 | 28.5 | 16.5 | 25.1 | 16.5 | 30.3 | 9.9 |
| Total | 24.7 | 16.8 | 18.5 | 12.6 | 17.4 | 14.6 | 20.2 | 10.3 |
| Total |  |  |  |  |  |  |  |  |
| Adults | 17.1 | 6.4 | 11.8 | 8.0 | 9.9 | 4.8 | 12.8 | 4.4 |
| Older Children | 21.6 | 13.7 | 15.7 | 11.8 | 11.7 | 9.4 | 16.4 | 7.5 |
| Young Children | 34.2 | 18.0 | 18.2 | 7.9 | 23.4 | 15.8 | 25.3 | 7.7 |
| Total | 22.8 | 14.1 | 14.5 | 9.3 | 13.9 | 11.3 | 17.1 | 8.0 |

*Note:* $n_{subjects} = 49$ ; $n_{observations} = 578$ . Values are marginalized over the variable Amplitude.

**Table 2.** WAIC model comparison.

|  | Model | WAIC | SE | WAIC weight |
| --- | --- | --- | --- | --- |
| Baseline | m.0 | 4409.9 | 54.6 | 0.00 |
| Additive | m.1 | 4349.8 | 61.9 | 0.08 |
| 2-way Interactions | m.2 | 4345.6 | 63.2 | 0.65 |
|  | m.3 | 4354.0 | 62.4 | 0.01 |
|  | m.4 | 4351.6 | 61.4 | 0.03 |
| All 2-way Interactions | m.5 | 4349.7 | 62.8 | 0.08 |
| 3-way Interactions | m.6 | 4348.5 | 60.8 | 0.15 |

*Note:* $n_{subjects} = 49; n_{observations} = 578$

**Table 3.**
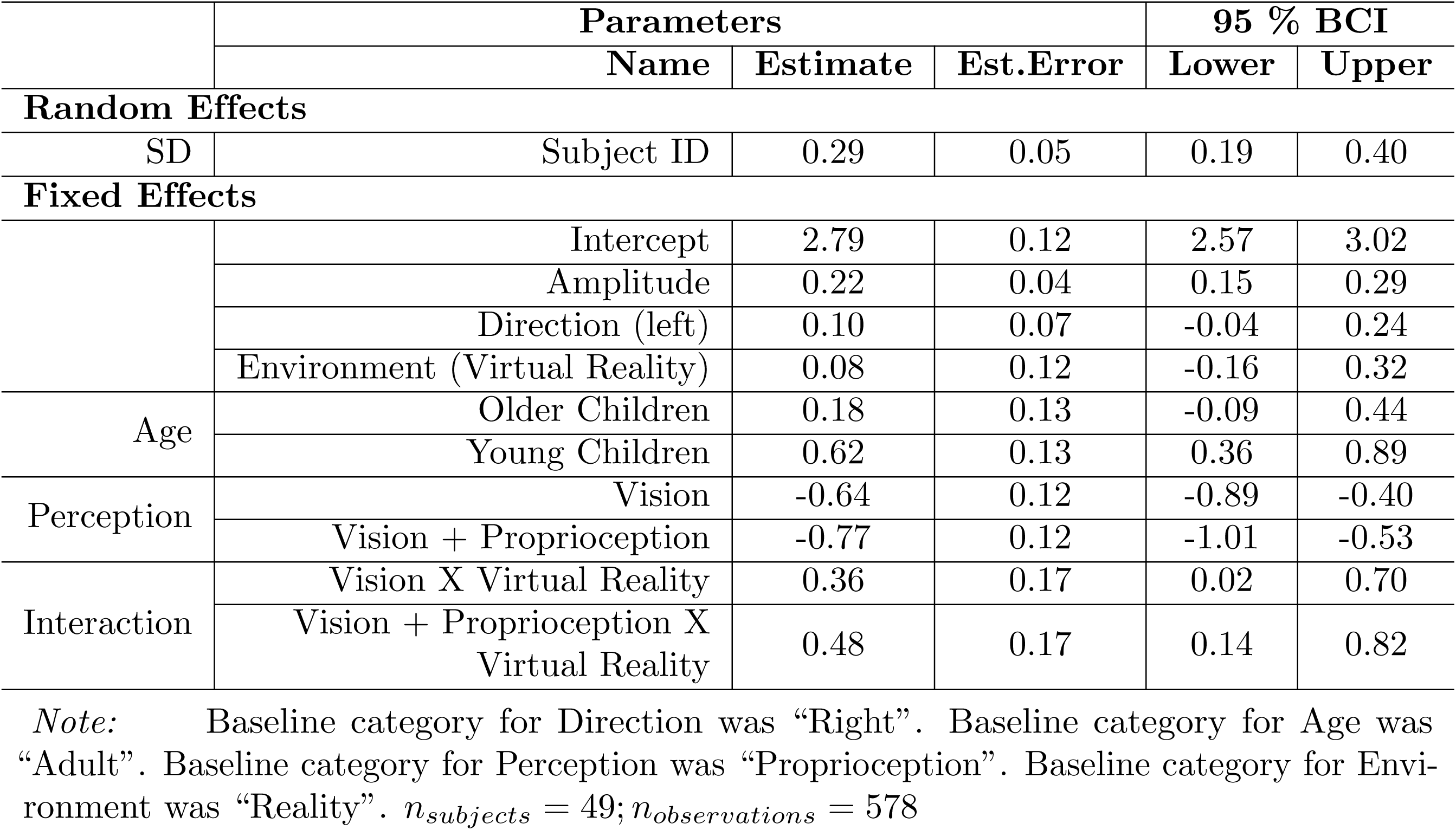
Estimated parameters of model m.2.

We considered Amplitude of the passive rotations as a continuous variable whose distribution is shown in Fig 3. To obtain interpretable results in the analyses, the Amplitude variable was standardized.

The mean self-turn error in the present sample was 17.1 degrees (SD = 8.0). The frequency of the observed values is reported in Fig 4. Considering how we computed the self-turn error, only positive values are possible and from visual inspection, the dependent variable has an evident positive skewed distribution.

**Fig 3.**
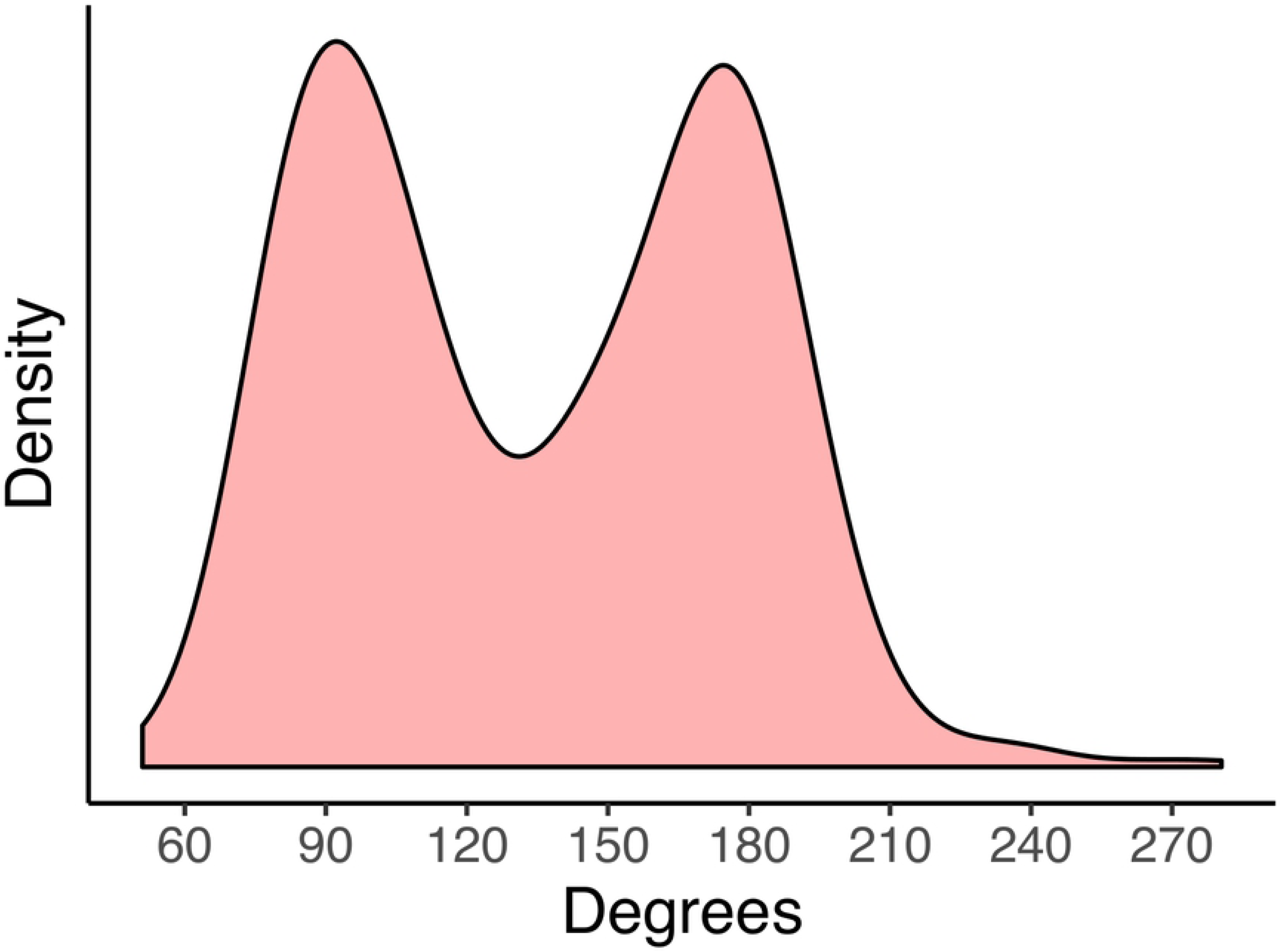
Estimated distribution of the actual Amplitude in the passive rotation. (*n*_*subjects*_ = 49; *n*_*observations*_ = 578)

The means and standard deviations of the self-turn error for the three age groups in the six different experimental conditions are reported in Table 1 and the distributions of the observed data are presented in Fig 5. For the sake of interpretability, descriptive statistics were marginalized over the variable Amplitude (integrating out this more imprecise variable) which will be considered later on in the analysis. Considering the marginal effect of Age, adults (M = 12.8, SD = 4.4) made less self-turn errors than older children (M = 16.4, SD = 7.5) and young children (M = 25.3, SD = 7.7). Looking at the marginal effect of Environment, subjects made less errors and were thusly more accurate in the reality condition (M = 13.9, SD = 8.0) than in the immersive virtual reality condition (M = 20.2, SD=10.3). Finally, for the marginal effect of Perception, subjects made less self-turn errors when they could rely on both vision and proprioception (M = 13.9, SD= 11.3) than when they could use only vision (M = 14.5, SD= 9.3) or proprioception (M = 22.8, SD= 14.1).

### Model comparison

Seven different Bayesian generalized mixed-effects models were performed to analyze the data (see Table 5 in S1 Supplemental Materials). In each model the dependent variable was the error in the self-turn task. The first model (m.0) was a baseline model considering the random effect of subjects (i.e. the random intercept that accounts for interpersonal variability) and the fixed effects of Direction (i.e. right or left rotation) and of Amplitude (i.e. amplitude of the rotation in degrees, reflecting the difficulty of the task). This baseline model, which includes the effects of possible confounding variables, was used as a reference point to then evaluate the models that considered the effects of Age, Perception, and Environment conditions. In the additive model (m.1) the additive effects of Age, Perception, and Environment were added to the baseline model. Single 2-way interactions were evaluated in models m.2, m.3, and m.4. These models respectively added to the additive model (m.1) the interaction effect between Perception and Environment conditions (m.2), Age and Environment conditions (m.3), and between Age and Perception (m.4). In the model m.5 all the possible 2-way interactions between Age, Perception, and Environment conditions were considered, together with the effects of the additive model (m.1). Finally, in the last model (m.6), the 3-way interaction between Age, Perception, and Environment conditions was added to the previous model (m.5).

**Table 4.** Predicted means and differences of Self-turn error according to Age.

| Mean |  | 95% BCI |  |
| --- | --- | --- | --- |
|  | Estimate | Lower | Upper |
| **Groups** |  |  |  |
| Adults | 12.8 | 10.6 | 15.1 |
| Older Children | 15.5 | 12.0 | 19.1 |
| Young Children | 24.8 | 19.3 | 30.7 |
| **Comparisons** |  |  |  |
| Young Children - Adult | 12.0 | 6.1 | 18.0 |
| Young - Older Children | 9.3 | 2.8 | 15.9 |
| Older Children - Adult | 2.7 | -1.4 | 6.8 |

*Note:* $n_{subjects} = 49; n_{observations} = 578$

**Table 5.**
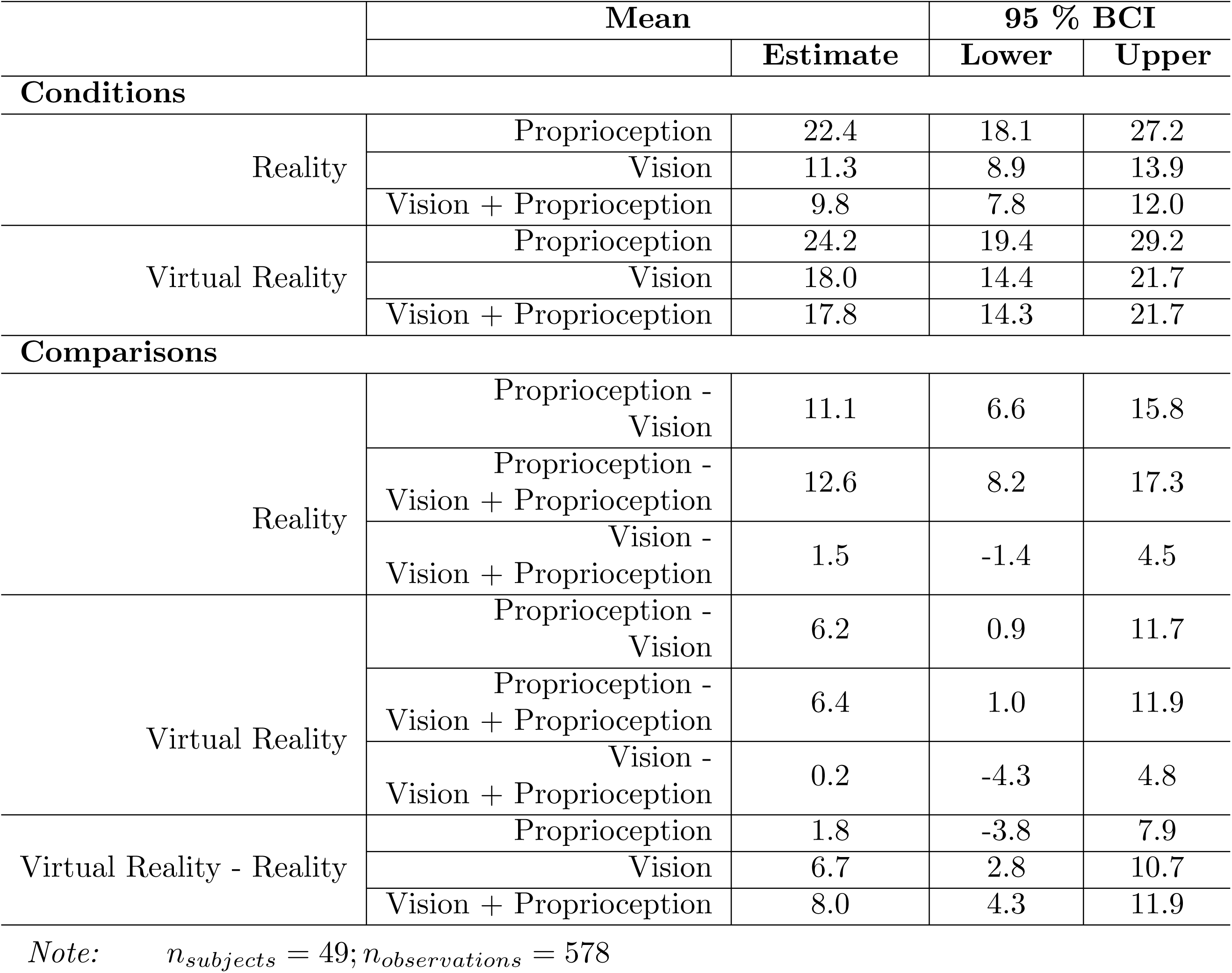
Predicted means and differences of Self-turn error according experimental conditions.

WAIC results indicated that m.2 was the most plausible model for the observed data, having the lower WAIC value (WAIC = 4345.6) and a probability of being the best of .65. Compared with the second-most plausible model (m.6 with a probability of .15), m.2 is 4.4 times more probable. WAIC values and relative WAIC weights or all models are reported in Table 2.

### Model interpretation

In order to interpret the effects of model m.2, 95% Bayesian Credible Intervals (BCIs) of the parameters posterior distribution were evaluated (Table 3). Ninety-five percent BCIs represent the range of the 95% most credible parameters values given the prior distribution and the observed data. Thus, an effect is considered to be present if the value zero is not included in the 95%BCI, whereas if the value zero is included in the 95%BCI, it is interpreted as no-effect.

Self-turn error was moderated by Amplitude, by Age, and by the interaction between Perception an Environment conditions. On the contrary, the direction of rotations seems to have no effect of the subjects’ performance (*β* = .10; 95% BCI = -.04; .24).

To evaluate the model fit (i.e. the model ability to explain the data) we used a Bayesian definition of R-squared [67] to estimate the proportion of variance explained. The estimated value of Bayesian R-squared for the model m.2 is .26 (95% BCI = .19; .34), that is the model explains 26% of the variability of the data.

#### Rotation amplitude

Self-turn error was moderated by Amplitude (*β* = .22; 95% BCI = .15; .29), for which increasing rotation amplitude is associated with a worse performance (Fig 6).

**Fig 4.**
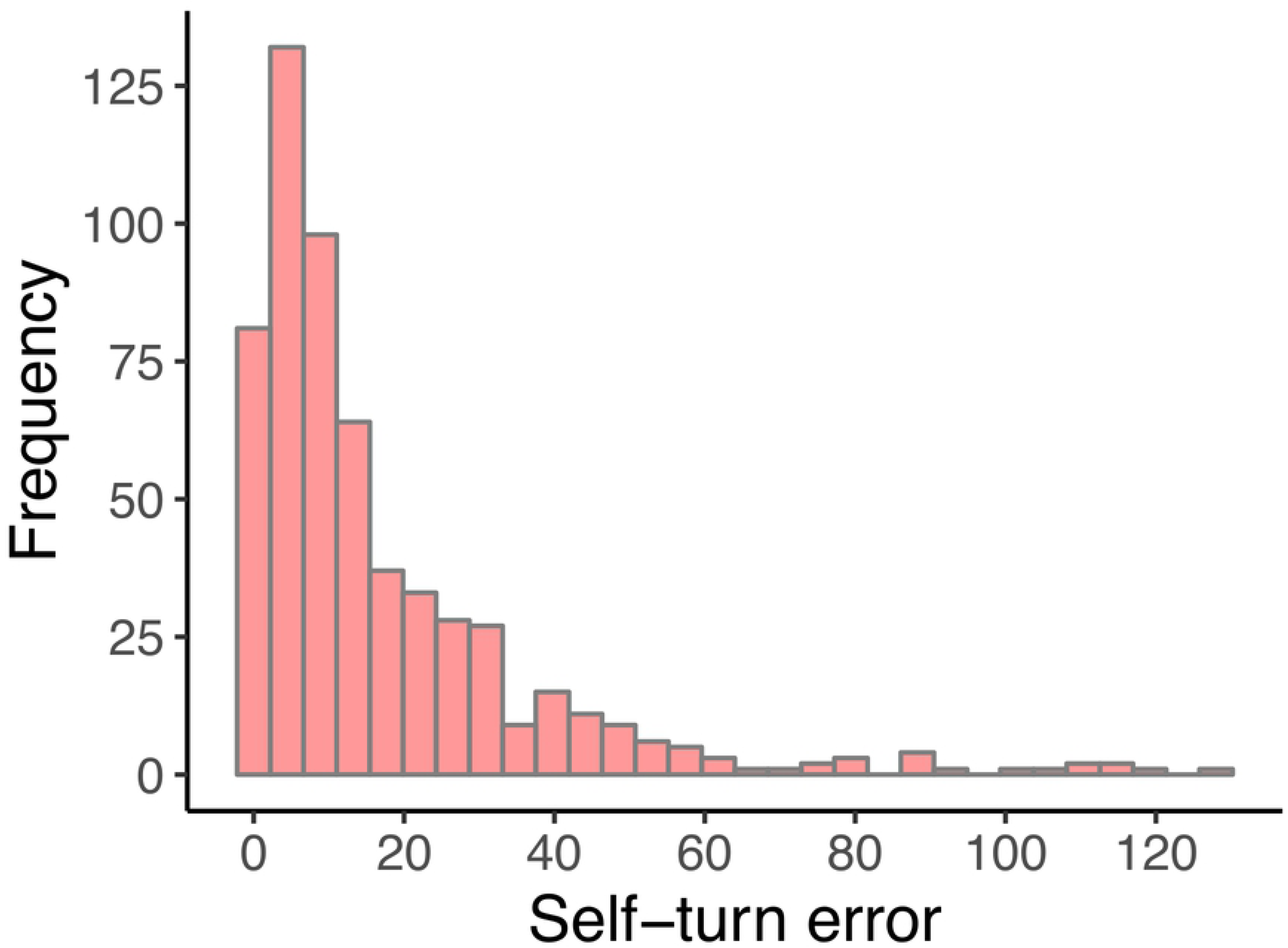
Frequencies of the observed self-turn errors. (*n*_*subjects*_ = 49; *n*_*observations*_ = 578)

**Fig 5.**
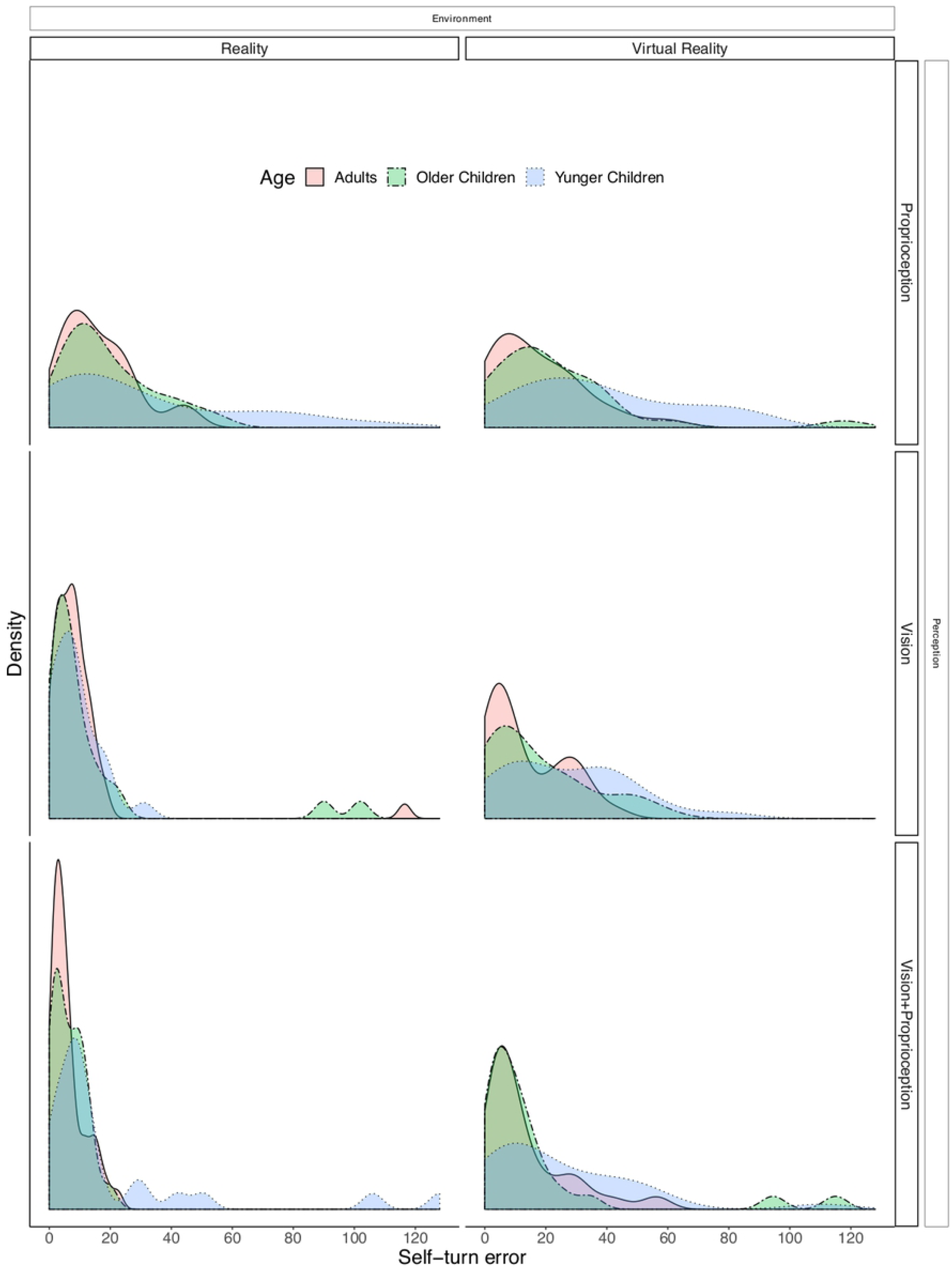
Estimated distributions of the observed self-turn errors in the different conditions according to age. (*n*_*subjects*_ = 49; *n*_*observations*_ = 578)

**Fig 6.**
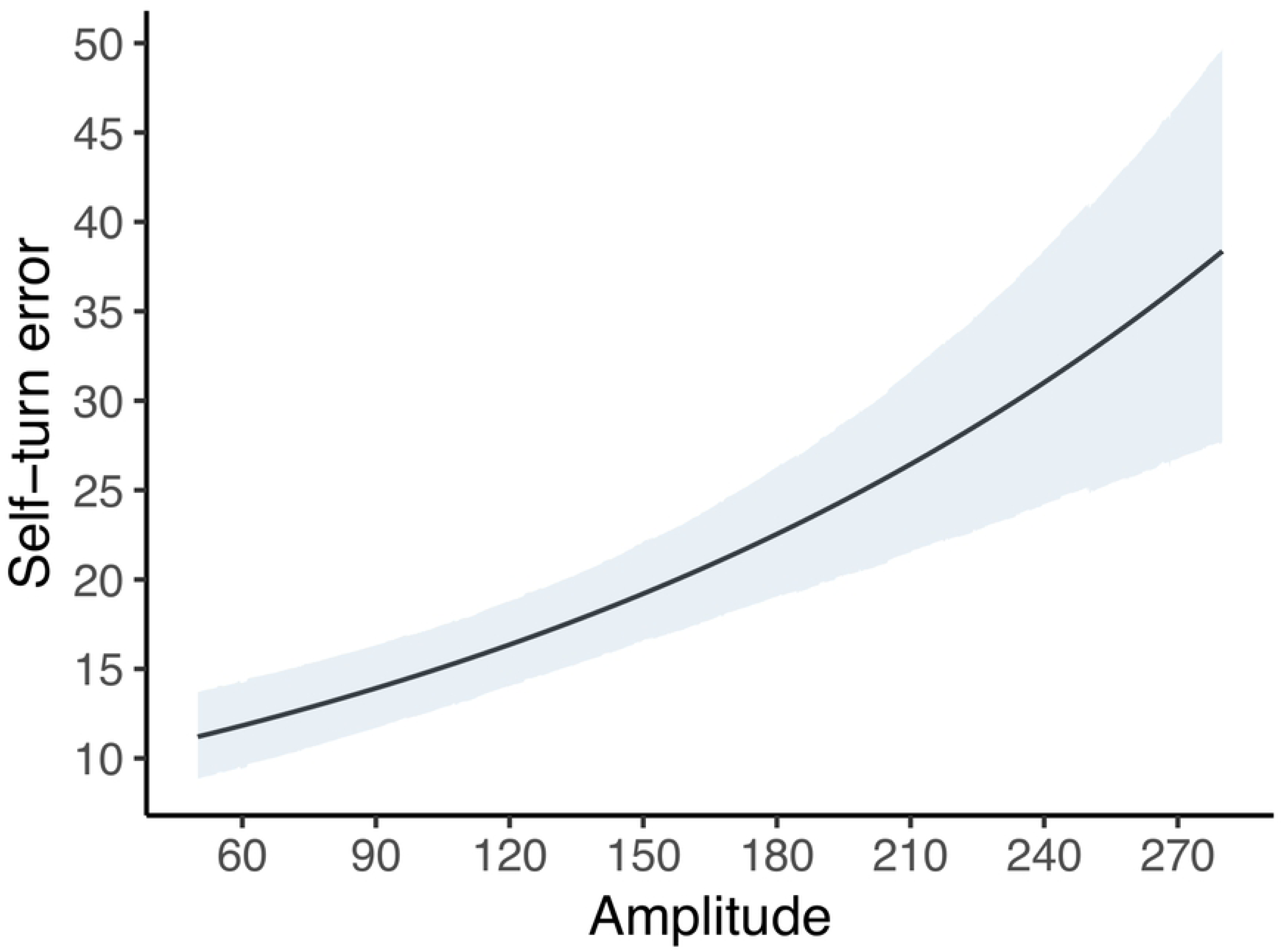
Predicted mean of self-turn error according to Amplitude. (*n*_*subjects*_ = 49; *n*_*observations*_ = 578). The line represents the mean value. The shaded area the 95% BCI values.

#### Group age

To evaluate the role of Age, the distributions of predicted mean values for the three groups were considered (Fig 7). The predicted mean error for adults was 12.8 degrees (95% BCI = 10.6;15.1), for older children 15.5 degrees (95% BCI = 12.0; 19.1) and for young children was 24.8 degrees (95% BCI = 19.3; 30.7). Bayesian pairwise comparisons (i.e. predicted score differences between groups) are reported in Table 4. Results showed that overall, young children are expected to make more self-turn errors than adults (95% BCI = 6.1; 18.0) and also more than older children (95% BCI = 2.8; 15.9). However, we cannot state that older children are expected to make more self-turn error because the 95% BCI of the difference includes the value zero (95% BCI = -1.4; 6.8).

**Fig 7.**
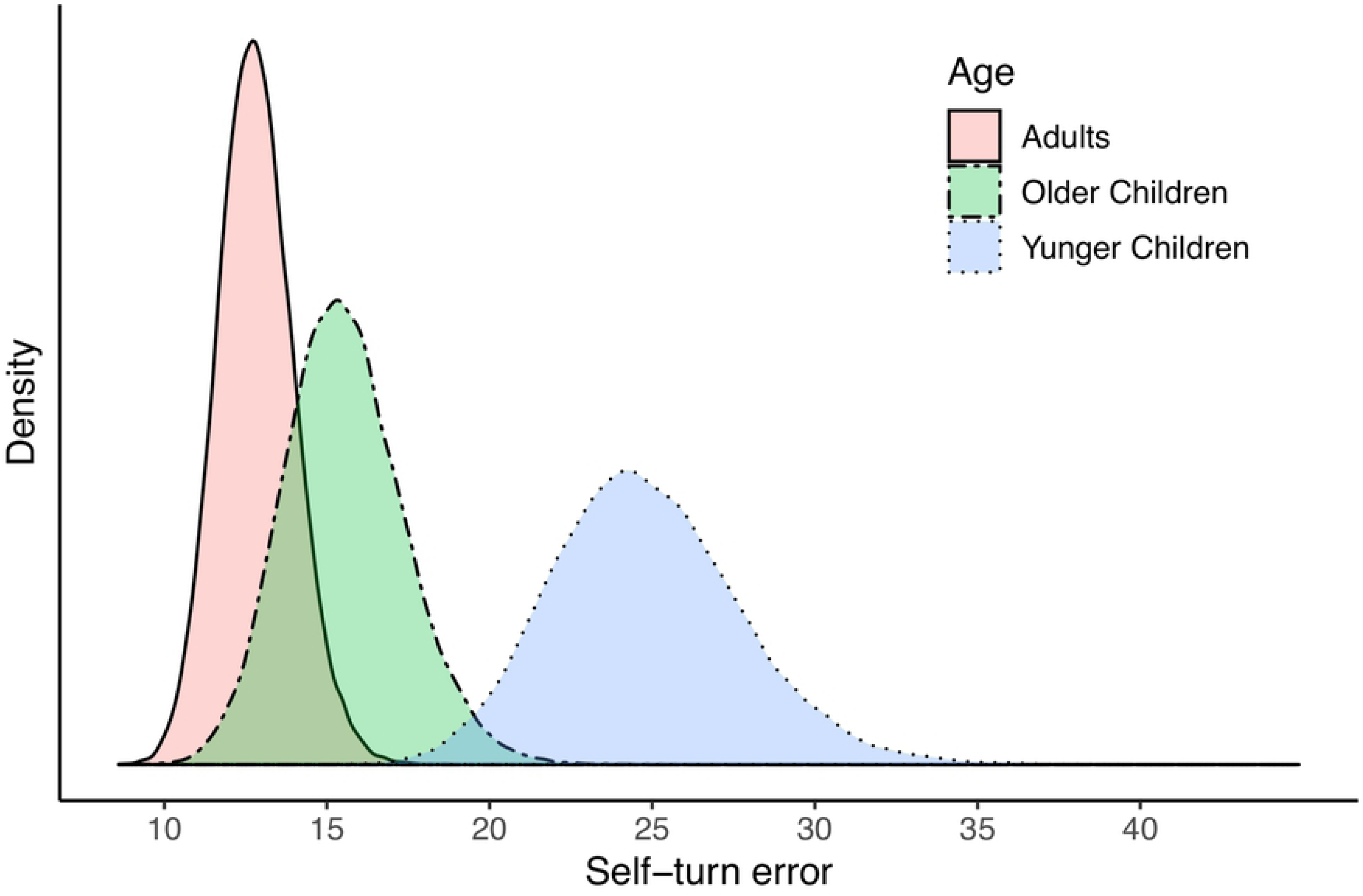
Distributions of the predicted means of self-turn error according to Age. (*n*_*subjects*_ = 49; *n*_*observations*_ = 578.

#### Perception and environment

To interpret the interaction between the Perception and Environment conditions, the distributions of predicted mean values for all six conditions were considered (Fig 8). In the Reality environment condition, the predicted mean error for proprioception was 22.4 degrees (95% BCI = 18.1; 27.2), for vision was 11.3 degrees (95% BCI = 8.9; 13.9) and for vision + proprioception was 9.8 degrees (95% BCI = 7.8; 12.0). In the Immersive Virtual Reality environment condition, the predicted mean error for proprioception was 22.4 degrees (95% BCI = 19.4; 29.2), for vision was 18.0 degrees (95% BCI = 14.4; 21.7) and for vision + proprioception was 17.8 degrees (95% BCI = 14.3; 21.7). Bayesian pairwise comparisons (i.e predicted error differences between conditions) are reported in Table 5. Results showed that in both Reality and Immersive Virtual Reality, subjects are expected to make more self-turn errors when they rely only on proprioception than when they can use only vision (Reality: 95% BCI = 6.6; 15.8; Immersive Virtual Reality: 95% BCI = 0.9; 11.7) or vision + proprioception (Reality: 95% BCI = 8.2; 17.3; Immersive Virtual Reality: 95% BCI = 1.0; 11.9). In addition, in both environments there is no difference between the use of vision and vision + proprioception (Reality: 95% BCI = -1.4; 4.5; Immersive Virtual Reality: 95% BCI = -4.3; 4.8). Moreover, comparing Immersive Virtual Reality to Reality conditions, results show that while wearing the HMD the self-turn errors increase when subjects rely only on vision (95% BCI = 2.8; 10.7) or on vision + proprioception (95% BCI = 4.3; 11.9), but subjects are not expected to make more errors than in Reality when they rely only on proprioception (95% BCI = -3.8; 7.9).

**Fig 8.**
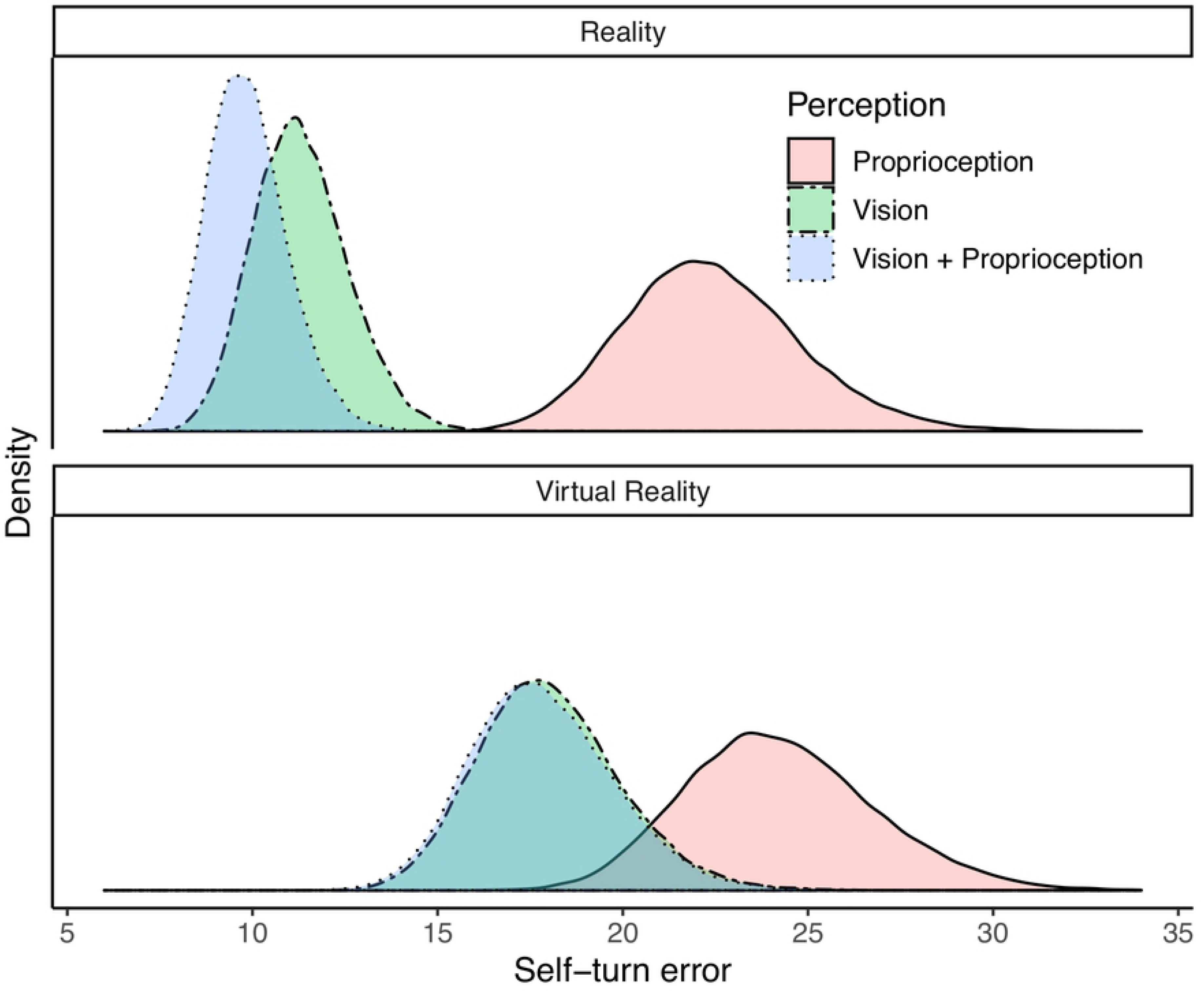
Distributions of the predicted means of self-turn error according to the different conditions. (*n*_*subjects*_ = 49; *n*_*observations*_ = 578)

#### Effect size

To quantify the differences between the various age groups and conditions, we expressed the effects as the ratio between the two scores of the comparison of interest (see Table 17 in S1 Supplemental Materials). Thus, for example, young children are expected to make 88% more errors than adults and 58% more errors than older children. Considering the Reality environment conditions, when using only proprioception subjects are expected to make 92% more errors than when they rely only on vision and 118% more errors than when using vision + proprioception. Considering the Immersive Virtual Reality environment conditions, when using only proprioception subjects are expected to make 34% more errors than when they rely only on vision and 35% more errors than when using vision + proprioception. Moreover, comparing Immersive Virtual Reality to Reality environmental condition, in IVR subjects are expected to make 56% more errors when using only vision and 75% when using vision + proprioception.

## Discussion

This experiment explored the extent to which visual information aids proprioceptive-based self-motion accuracy across the lifespan, and specifically in three developmental groups: 4–8-year-old children, 9–15-year-old children, and adults. Moreover, the experiment assessed whether HMD-delivered IVR environments affect accuracy.

As expected, we found a main developmental trend in the improvement of proprioception across conditions. In particular, as hypothesized, we found differences between the young child group (4–8 years old) and the older child and adult groups (9–15 and 20–43 years old), with this youngest group showing lower proprioceptive accuracy than the two older groups. This indicates that proprioceptive development predominantly takes place in the first eight years of life, such that adolescent and pre-adolescent children make more accurate proprioceptive judgements than younger children.

In line with our hypotheses, we also found an interaction effect between Perception and Environment condition. Our findings indicate that proprioceptive accuracy was markedly impaired when participants could rely only on proprioceptive input, regardless of the environment. In the conditions which forced participants to rely solely on proprioception by removing all visual information, all groups were less accurate than in conditions where visual information was provided, regardless of the salience of this visual information. This finding is consistent with the assertion that visual and vestibular information combine with proprioceptive information to allow accurate self-motion [11]. Moreover, it indicates that typically developing child and adult populations rely specifically on vision to calibrate proprioception in order to accurately judge their movements. Regarding the role of different visual landmarks, no differences were found between vision + proprioception and vision only conditions, that is, conditions in which participants could view all aspects of the real or virtual room versus conditions in which participants received visual input of randomly placed clouds but were unable to see visual landmarks such as the corners of the room or their body. Moreover, IVR, compared to Reality, disrupted proprioception only when visual input was provided (vision + proprioception and vision only conditions). There were no differences between IVR vs Reality in only proprioception (blind) conditions. This allows us to exclude the possibility that wearing the HMD alone, and the corresponding weight and head restriction, might have disrupted proprioception. We did find that performance worsened in IVR conditions where visual information was available relative to corresponding reality conditions. The way in which the HMD delivers visual information has a complex (and essentially unknown) effect on self-motion perception and the kinematics of movement [68]. Factors such as display type, field of view, visual content (peripheral cues, high-low visual contrast, etc.), temporal lag between the user’s action and the HMD’s reaction, and so on could be the means by which IVR disrupts proprioception through vision. This is an important finding, given that few IVR experiments have considered that performance may be affected simply due to the use of IVR or HMD-delivered IVR. Many previous IVR experiments seem to implicitly assume that performance in IVR constitutes an appropriate corollary for real-world performance, but our findings indicate that this may not be the case. Despite this HMD effect, our results provides evidence that IVR may be a useful means of studying multisensory integration and accuracy. Indeed, the same general Perception trend in self-motion accuracy (proprioception only¡vision only vision + proprioception) was found both in IVR and R environments.

In contrast to our expectations, we failed to find any Age x Condition interaction effect. We expected that adults would be more affected by disrupted proprioception than children, but this was not the case. Various aspects of the experimental design should be taken into account to discuss this result. Firstly, our manipulation of the multisensory input in different conditions could have been insufficient to uncover the expected differences. We found the expected general trend of reduced proprioceptive accuracy in vision conditions relative to vision + proprioception conditions. However, this difference failed to reach meaningful magnitude. As previous studies highlight, relative dominance of visual and proprioceptive input and visuo-proprioceptive integration are task-dependent [2, 30]. For example, proprioception was reported to be more precise in the radial (near-far) direction and vision in the azimuthal (left-right) direction [69–71]. It could be suggested that our azimuthal proprioceptive task was too dependent on vision to allow the detection of differences that were due to the disruption of proprioception. In fact, our vision conditions were designed to disrupt proprioception by removing visually-driven proprioceptive information (the room corners and participant’s body), while still providing non-proprioceptive visual landmarks (surrounding clouds). It could be the case that non-proprioceptively salient visual landmarks are sufficient to allow accurate performance in our task. In addition, other similar studies used a standing self-turn paradigm [53, 72]. We utilized a seated self-turn paradigm so that we could use the chair position as a measurement point of reference, independently from the participants’ individual postures which may vary. The sitting self-turn paradigm keeps the subject’s position in the center of the room, allowing us to make the task and measurement consistent across participants. However, this seated task could be less challenging than a standing one, resulting in a ceiling effect, particularly for older children and adult groups.

We also found a main effect of rotation amplitude, with proprioceptive accuracy consistently decreasing as rotation amplitude increased across conditions and groups. Despite the fact that studying the effect of rotation amplitude was not a primary goal of this work, it is interesting to speculate whether this effect may be specifically due to working memory constraints in larger rotations. Body position-matching tasks similar to the one used in the present study imply the need for executive skills. Indeed, current tests for the assessment of proprioception evaluate the reproduction of body positions or movements by relying on active rehearsal in working memory [49, 50]. In our experiment, accuracy largely depends on the ability of participants to actively maintain the start position in memory, and it may be the case that differences in working memory capacity across age groups could have affected results. Indeed, working memory limitations have been found up to pre-adolescence [73] and age-related lower visuo-spatial working memory capacity can be associated with lower proprioceptive accuracy in body position-matching tasks [74].

The present study opens intriguing perspectives for future research, despite having some limitations. Firstly, the experimenter manually rotated the participant, so although experimenters were trained to keep a similar speed and method of rotating, the rotation velocity was not perfectly consistent across trials and participants, potentially influencing participants’ performance as in previous works [72]. Another limitation concerned the manipulation of visual conditions distinguishing between “only vision” and “vision + proprioception”. As we found no meaningful differences between these two Perception conditions, the “only vision” condition could have been insufficient to isolate vision and disrupt proprioception as we aimed to. It would be interesting to see how similar but more effective manipulations of visual information aimed at disrupting proprioception would affect performance compared to conditions where instead only optic flow is available (i.e. no movement). As previously mentioned, it is also possible that self-motion differences in these two conditions were too small to be detected with our task, and might be elicited with a more difficult one. Moreover, the age groups could be too broad to clearly show early developmental trends and changes. Further research could focus specifically on children younger than eight years old to explore the early development of visuo-proprioceptive integration. Furthermore, future studies could utilize our paradigm to explore age-related visuo-spatial working memory abilities associated with proprioception. A more in-depth look is also necessary to investigate potential implications of a proprioceptive sensory register and its influence on performance in multisensory motor tasks, as individual sensory registers have been shown to affect working memory in multisensory environments (for a review, see [75]).

One of the most intriguing yet unexplored perspectives that led to this work concerns the possibility of intentionally disrupting proprioception through HMD-delivered IVRs. This method could be employed to study the degree to which different developmental populations rely on proprioception, vision, and visuo-proprioceptive integration. From an applied perspective, disrupting proprioception could comprise an innovative intervention for use with clinical populations which demonstrate an atypical reliance on specific senses and atypical integration of vision (*exteroception*) and proprioception. For example, people with Autism Spectrum Disorder (ASD) seem to show an over-reliance on proprioception and hypo-reliance on exteroception [76–78]. This perceptual strategy might not only lead to impaired motor skills in ASD (e.g. dyspraxia and repetitive behaviors), but also seems to be related to core features of impaired social and communicative development. Interventions could be aimed at increasing the reliance on vision in children with ASD by disrupting proprioception. In this respect a possible speculation is that IVR interventions could constitute a useful training method to achieve a therapeutic purpose.

## Conclusion

In sum, the present study offers useful insights regarding the use of IVR in research on multisensory integration and sensorimotor functioning. When visual information is provided, proprioceptive accuracy in IVR seems to be impaired relative to performance in reality. As proprioception is fundamental to performance in any motor task, this has to be taken into account when interpreting the results of IVR studies which involve proprioceptive abilities. However, IVR could still be a useful tool for detecting multisensory trends. In fact, we found the same condition-specific trend in IVR as in reality. Both in reality and IVR, the conditions which allowed a reliance solely on proprioception led to the lowest proprioceptive accuracy, and minimal differences emerged between vision only and vision + proprioception conditions. The exploratory nature of the present study could contribute to the undertaking of more confirmatory future studies, which would benefit from the estimated effect sizes provided here, to develop and test further hypotheses.

## Aknowledgments

Our gratitude to the multimedia designers Marco Godeas and Carlo Marzaroli. They did a great job in building the IVR environment and setting up the laboratory. This research was helped immensely by their experience with IVR technologies and children with Autism Spectrum Disorder. This experiment would not have been possible without their ideas and technical support.

Thanks to F.lli Budai S.r.l. for building the experimental room: a generous gift for which we are very grateful.

Thanks to Beneficentia Stiftung Foundation for supporting our research.

Sincere thanks to Associazione Pro Musica Ruda – Scuola Comunale di Musica and the Comune di Ruda (Udine, Italy) for hosting our laboratory and supporting our work.

## Author Contributions

**Conceptualization:** Irene Valori, Phoebe McKenna-Plumley, Rena Bayramova, Teresa Farroni.

**Data Curation:** Claudio Zandonella Callegher, Irene Valori, Phoebe McKenna-Plumley, Rena Bayramova.

**Formal Analysis:** Claudio Zandonella Callegher, Gianmarco Altoè, Irene Valori.

**Funding Acquisition:** Teresa Farroni.

**Investigation:** Irene Valori, Phoebe McKenna-Plumley, Rena Bayramova, Teresa Farroni.

**Methodology:** Irene Valori, Phoebe McKenna-Plumley, Rena Bayramova, Teresa Farroni.

**Project Administration:** Irene Valori, Teresa Farroni.

**Resources:** Teresa Farroni.

**Supervision:** Teresa Farroni.

**Visualization:** Claudio Zandonella Callegher, Irene Valori, Phoebe McKenna-Plumley, Rena Bayramova, Gianmarco Altoé.

**Writing – Original Draft Preparation:** Irene Valori, Phoebe McKenna-Plumley, Rena Bayramova, Claudio Zandonella Callegher, Teresa Farroni.

**Writing – Review & Editing:** Phoebe McKenna-Plumley, Irene Valori, Rena Bayramova, Claudio Zandonella Callegher, Gianmarco Altoé, Teresa Farroni.

## References

1. Craig AD. How Do You Feel? Interoception: The Sense of the Physiological Condition of the Body. Nature Reviews Neuroscience. 2002;3(8):655–666. doi:10.1038/nrn894.

2. Bremner AJ, Lewkowicz DJ, Spence C. The Multisensory Approach to Development. In: Bremner AJ, Lewkowicz DJ, Spence C, editors. Multisensory Development. Oxford: United Kingdom: Oxford University Press; 2012. p. 1–26.

3. Knudsen EI. Instructed Learning in the Auditory Localization Pathway of the Barn Owl. Nature. 2002;417(6886):322–328. doi:10.1038/417322a.

4. Scarfe P, Hibbard PB. Statistically Optimal Integration of Biased Sensory Estimates. Journal of Vision. 2011;11(7):12. doi:10.1167/11.7.12.

5. Spence C. Explaining the Colavita Visual Dominance Effect. Progress in Brain Research. 2009;176:245–258. doi:10.1016/S0079-6123(09)17615-X.

6. Nelson RJ. The Somatosensory System: Deciphering the Brain’s Own Body Image. 1st ed. Boca Raton, Florida: CRC Press; 2001.

7. Stillman BC. Making Sense of Proprioception: The Meaning of Proprioception, Kinaesthesia and Related Terms. Physiotherapy. 2002;88(11):667–676. doi:10.1016/S0031-9406(05)60109-5.

8. Pereira EM, Rueda FM, Diego IA, De La Cuerda RC, De Mauro A, Page JM. Use of Virtual Reality Systems as Proprioception Method in Cerebral Palsy: Clinical Practice Guideline. Neurología (English Edition). 2014;29(9):550–559. doi:10.1016/j.nrleng.2011.12.011.

9. Ramachandran VS, Rogers-Ramachandran D. Synaesthesia in Phantom Limbs Induced with Mirrors. Proceedings of the Royal Society B. 1996;263(1369):377–386. doi:10.1098/rspb.1996.0058.

10. Tsakiris M, Hesse MD, Boy C, Haggard P, Fink GR. Neural Signatures of Body Ownership: A Sensory Network for Bodily Self-Consciousness. Cerebral Cortex. 2006;17(10):2235–2244. doi:10.1093/cercor/bhl131.

11. Cullen KE. The Vestibular System: Multimodal Integration and Encoding of Self-Motion for Motor Control. Trends in Neurosciences. 2012;35(3):185–196.

12. Bremner AJ, Holmes NP, Spence C. The Development of Multisensory Representations of the Body and of the Space around the Body. In: Bremner AJ, Lewkowicz DJ, Spence C, editors. Multisensory Development. Oxford: United Kingdom: Oxford University Press; 2012. p. 113–136.

13. Sigmundsson H, Whiting HTA, Loftesnes JM. Development of Proprioceptive Sensitivity. Experimental Brain Research. 2000;135(3):348–352. doi:10.1007/s002210000531.

14. von Hofsten C, Rösblad B. The Integration of Sensory Information in the Development of Precise Manual Pointing. Neuropsychologia. 1988;26(6):805–821. doi:10.1016/0028-3932(88)90051-6.

15. Goble DJ, Lewis CA, Hurvitz EA, Brown SH. Development of Upper Limb Proprioceptive Accuracy in Children and Adolescents. Human Movement Science. 2005;24(2):155–170. doi:10.1016/j.humov.2005.05.004.

16. Hearn M, Crowe A, Keessen W. Influence of Age on Proprioceptive Accuracy in Two Dimensions. Perceptual and Motor Skills. 1989;69(3–1):811–818. doi:10.1177/00315125890693-118.

17. Nardini M, Cowie D. The Development of Multisensory Balance, Locomotion, Orientation and Navigation. In: Bremner AJ, Lewkowicz DJ, Spence C, editors. Multisensory Development. Oxford: United Kingdom: Oxford University Press; 2012. p. 137–158.

18. Bremner AJ, Hill EL, Pratt M, Rigato S, Spence C. Bodily Illusions in Young Children: Developmental Change in Visual and Proprioceptive Contributions to Perceived Hand Position. PLoS ONE. 2013;8(1):51887. doi:10.1371/journal.pone.0051887.

19. Sanchez-Vives MV, Slater M. From Presence to Consciousness through Virtual Reality. Nature Reviews Neuroscience. 2005;6(4):332–339. doi:10.1038/nrn1651.

20. Gromala D, Shaw C, Song M. Chronic Pain and the Modulation of Self in Immersive Virtual Reality. In: Chair AVS, editor. Biologically Inspired Cognitive Architectures, Papers from the 2009 AAAI Fall Symposium, Arlington, Virginia, USA, November 5–7, 2009. Symposium Conducted at the 2009 AAAI Fall Symposium Series, Arlington, VA.; 2009. p. 71.

21. Murray CD, Sixsmith J. The Corporeal Body in Virtual Reality. Ethos. 1999;27(3):315–343. doi:10.1525/eth.1999.27.3.315.

22. Mohler BJ, Campos JL, Weyel M, Bülthoff HH. Gait Parameters While Walking in a Head-Mounted Display Virtual Environment and the Real World. In: Fröhlich B, Blach R, van Liere R, editors. 13th Eurographics Symposium on Virtual Environments and 10th Immersive Projection Technology Workshop (IPT-EGVE 2007). Weimar, Germany: Eurographics Association; 2007. p. 1–4.

23. Riecke BE, Wiener JM. Can People Not Tell Left from Right in VR? Point-to-Origin Studies Revealed Qualitative Errors in Visual Path Integration. In: 2007 IEEE Virtual Reality Conference, Charlotte, NC. Piscataway, NJ: IEEE; 2007. p. 3–10.

24. Prothero JD, Parker DE. A Unified Approach to Presence and Motion Sickness. In: Hettinger L, and MH, editors. Virtual and Adaptive Environments: Applications, Implications, and Human Performance Issues. Mahwah, NJ: Lawrence Erlbaum Associates Publishers; 2003. p. 47–66.

25. Campos JL, H BH. Multimodal Integration during Self-Motion in Virtual Reality. In: Murray MM, Wallace MT, editors. The Neural Bases of Multisensory Processes. Boca Raton, Florida: CRC Press/Taylor & Francis; 2012. p. 603–628.

26. Riecke BE, Schulte-Pelkum J, Buelthoff HH. Perceiving Simulated Ego-Motions in Virtual Reality: Comparing Large Screen Displays with HMDs. In: SPIE 2005; 2005. p. 344–356.

27. Kearns MJ, Warren WH, Duchon AP, Tarr MJ. Path Integration from Optic Flow and Body Senses in a Homing Task. Perception. 2002;31(3):349–374. doi:10.1068/p3311.

28. Bakker NH, Werkhoven PJ, Passenier PO. The Effects of Proprioceptive and Visual Feedback on Geographical Orientation in Virtual Environments. Presence: Teleoperators & Virtual Environments. 1999;8(1):36–53. doi:10.1162/105474699566035.

29. Lathrop WB, Kaiser MK. Perceived Orientation in Physical and Virtual Environments: Changes in Perceived Orientation as a Function of Idiothetic Information Available. Presence: Teleoperators & Virtual Environments. 2002;11(1):19–32. doi:10.1162/105474602317343631.

30. Adams H, Narasimham G, Rieser J, Creem-Regehr S, Stefanucci J, Bodenheimer B. Locomotive Recalibration and Prism Adaptation of Children and Teens in Immersive Virtual Environments. IEEE Transactions on Visualization & Computer Graphics. 2018;24(4):1408–1417. doi:10.1109/TVCG.2018.2794072.

31. Bodenheimer B, Creem-Regehr S, Stefanucci J, Shemetova E, Thompson WB. Prism Aftereffects for Throwing with a Self-Avatar in an Immersive Virtual Environment. In: Rosenberg ES, editor. 2017 IEEE Virtual Reality (VR) Proceedings: March 18-22, 2017, Los Angeles, CA, USA. Piscataway, NJ: IEEE; 2017. p. 141–147.

32. Mohler BJ, Thompson WB, Creem-Regehr SH, Willemsen P, Pick Jr HL, Rieser JJ. Calibration of Locomotion Resulting from Visual Motion in a Treadmill-Based Virtual Environment. ACM Transactions on Applied Perception (TAP). 2007;4(1):1–15. doi:10.1145/1227134.1227138.

33. Petrini K, Caradonna A, Foster C, Burgess N, Nardini M. How Vision and Self-Motion Combine or Compete during Path Reproduction Changes with Age. Scientific Reports. 2016;6:29163. doi:10.1038/srep29163.

34. Petrini K, Jones PR, Smith L, Nardini M. Hearing Where the Eyes See: Children Use an Irrelevant Visual Cue When Localizing Sounds. Child Development. 2015;86:1449–1457. doi:10.1111/cdev.12397.

35. Bailey JO, Bailenson JN. Immersive Virtual Reality and the Developing Child. In: Blumberg FC, Brooks PJ, editors. Cognitive Development in Digital Contexts. Amsterdam, Netherlands: Elsevier; 2017. p. 181–200.

36. McElreath R. Statistical Rethinking: A Bayesian Course with Examples in R. New York, NY: Chapman and Hall/CRC; 2015.

37. Wagenmakers EJ, Farrell S. AIC Model Selection Using Akaike Weights. Psychonomic Bulletin & Review. 2004;11(1):192–196. doi:10.3758/BF03206482.

38. Vandekerckhove J, Matzke D, Wagenmakers EJ. Model Comparison and the Principle of Parsimony. In: Busemeyer JR, Wang Z, Townsend JT, Eidels A, editors. The Oxford handbook of computational and mathematical psychology. New York, NY: Oxford University Press; 2015. p. 300–319.

39. Gelman A, Stern HS, Carlin JB, Dunson DB, Vehtari A, Rubin DB. Bayesian Data Analysis. New York, NY: Chapman and Hall/CRC; 2013.

40. Kruschke JK, Liddell TM. The Bayesian New Statistics: Hypothesis Testing, Estimation, Meta-Analysis, and Power Analysis from a Bayesian Perspective. Psychonomic Bulletin & Review. 2018;25(1):178–206. doi:10.3758/s13423-016-1221-4.

41. Bolker BM, Brooks ME, Clark CJ, Geange SW, Poulsen JR, Stevens MHH, et al. Generalized Linear Mixed Models: A Practical Guide for Ecology and Evolution. Trends in Ecology & Evolution. 2009;24(3):127–135. doi:10.1016/j.tree.2008.10.008.

42. Fong Y, Rue H, Wakefield J. Bayesian Inference for Generalized Linear Mixed Models. Biostatistics. 2010;11(3):397–412. doi:10.1093/biostatistics/kxp053.

43. Romeijn JW, van de Schoot R. A Philosopher’s View on Bayesian Evaluation of Informative Hypotheses. In: Hoijtink H, Klugkist I, Boelen PA, editors. Bayesian Evaluation of Informative Hypotheses. New York, NY: Springer; 2008. p. 329–357.

44. Etz A, Vandekerckhove J. Introduction to Bayesian Inference for Psychology. Psychonomic Bulletin & Review. 2018;25(1):5–34. doi:10.3758/s13423-017-1262-3.

45. van de Schoot R, Kaplan D, Denissen J, Asendorpf JB, Neyer FJ, van Aken MAG. A Gentle Introduction to Bayesian Analysis: Applications to Developmental Research. Child Development. 2014;85(3):842–860. doi:10.1111/cdev.12169.

46. Hurley MV, Rees J, Newham DJ. Quadriceps Function, Proprioceptive Acuity and Functional Performance in Healthy Young, Middle-Aged and Elderly Subjects. Age and Ageing. 1998;27(1):55–62. doi:10.1093/ageing/27.1.55.

47. Wingert JR, Welder C, Foo P. Age-Related Hip Proprioception Declines: Effects on Postural Sway and Dynamic Balance. Archives of Physical Medicine and Rehabilitation. 2014;95(2):253–261. doi:10.1016/j.apmr.2013.08.012.

48. Jürgens R, Boss T, Becker W. Estimation of Self-Turning in the Dark: Comparison between Active and Passive Rotation. Experimental Brain Research. 1999;128(4):491–504. doi:10.1007/s002210050872.

49. Waddington G, Adams R. Discrimination of Active Plantarflexion and Inversion Movements after Ankle Injury. Journal of Physiotherapy. 1999;45(1):7–13. doi:10.1016/S0004-9514(14)60335-4.

50. Weerakkody NS, Blouin JS, Taylor JL, Gandevia SC. Local Subcutaneous and Muscle Pain Impairs Detection of Passive Movements at the Human Thumb. The Journal of Physiology. 2008;586(13):3183–3193. doi:10.1113/jphysiol.2008.152942.

51. Chen X, Treleaven J. The Effect of Neck Torsion on Joint Position Error in Subjects with Chronic Neck Pain. Manual Therapy. 2013;18(6):562–567. doi:10.1016/j.math.2013.05.015.

52. Hallgren KA. Computing Inter-Rater Reliability for Observational Data: An Overview and Tutorial. Tutorials in Quantitative Methods For. Psychology. 2012;8(1):23. doi:10.20982/tqmp.08.1.p023.

53. Wang RF, Spelke ES. Updating egocentric representations in human navigation. Cognition. 2000;77(3):215–250. doi:10.1016/S0010-0277(00)00105-0.

54. Meilinger T, Schulte-Pelkum J, Frankenstein J, Berger DR, Bülthoff HH. Global Landmarks Do Not Necessarily Improve Spatial Performance in Addition to Bodily Self-Movement Cues When Learning a Large-Scale Virtual Environment. In: Imura M, Figueroa P, Mohler B, editors. 25th International Conference on Artificial Reality and Telexistence and the 20th Eurographics Symposium on Virtual Environments, (ICAT-EGVE 2015), Kyoto, Japan. Aire-la-Ville, Switzerland: Eurographics Association; 2015. p. 25–28.

55. Pinheiro J, Bates D. Mixed-effects models in S and S-PLUS. New York, NY: Springer-Verlag; 2006.

56. Ng VKY, Cribbie RA. Using the Gamma Generalized Linear Model for Modeling Continuous, Skewed and Heteroscedastic Outcomes in Psychology. Current Psychology. 2017;36(2):225–235. doi:10.1007/s12144-015-9404-0.

57. R Core Team. R: A Language and Environment for Statistical Computing.; 2018. R Foundation for Statistical Computing.

58. Bürkner PC. brms: An R package for Bayesian multilevel models using Stan. Journal of Statistical Software. 2017;80(1):1–28. doi:10.18637/jss.v080.i01.

59. Stan Development Team. Stan: A C++ Library for Probability and Sampling.; 2017.

60. Stan Development Team. Stan Modeling Language: User’s Guide and Reference Manual; 2017.

61. Hoffman MD, Gelman A. The No-U-Turn Sampler: Adaptively Setting Path Lengths in Hamiltonian Monte Carlo. Journal of Machine Learning Research. 2014;15:1593–1623.

62. Neal RM. MCMC Using Hamiltonian Dynamics. In: Brooks S, Gelman A, Jones GL, Meng XL, editors. Handbook of Markov Chain Monte Carlo. New York, NY: Chapman; 2012. p. 113–162.

63. Gelman A, Rubin DB. Inference from Iterative Simulation Using Multiple Sequences. Statistical Science. 1992;7(4):457–472. doi:10.1214/ss/1177011136.

64. Gelman A, Hwang J, Vehtari A. Understanding Predictive Information Criteria for Bayesian Models. Statistics and Computing. 2014;24(6):997–1016. doi:10.1007/s11222-013-9416-2.

65. Vehtari A, Gelman A, Gabry J. Practical Bayesian Model Evaluation Using Leave-One-out Cross-Validation And. WAIC Statistics and Computing. 2017;27(5):1413–1432. doi:10.1007/s11222-016-9696-4.

66. Akaike H. Information Theory and an Extension of the Maximum Likelihood Principle. In: Petrov BN, Csaki F, editors. Proceedings of the Second International Symposium on Information Theory. Budapest Akademiai Kiado; 1973. p. 267–281.

67. Gelman A, Goodrich B, Gabry J, Vehtari A. R-Squared for Bayesian Regression Models. The American Statistician. 2018;1(6):10–1080. doi:10.1080/00031305.2018.1549100.

68. Powell WA, Stevens B. The Influence of Virtual Reality Systems on Walking Behaviour: A Toolset to Support Application Design. In: 2013 International Conference on Virtual Rehabilitation (ICVR), Philadelphia, PA. Piscataway, NJ: IEEE; 2013. p. 270–276.

69. Snijders HJ, Holmes NP, Spence C. Direction-Dependent Integration of Vision and Proprioception in Reaching under the Influence of the Mirror Illusion. Neuropsychologia. 2007;45(3):496–505. doi:10.1016/j.neuropsychologia.2006.01.003.

70. van Beers RJ, Sittig AC, van der Gon DJJ. Integration of proprioceptive and visual position-information: An experimentally supported model. Journal of Neurophysiology. 1999;81(3):1355–1364. doi:10.1152/jn.1999.81.3.1355.

71. van Beers RJ, Wolpert DM, Haggard P. When Feeling Is More Important than Seeing in Sensorimotor Adaptation. Current Biology. 2002;12(10):834–837. doi:10.1016/S0960-9822(02)00836-9.

72. Jürgens R, Becker W. Perception of Angular Displacement without Landmarks: Evidence for Bayesian Fusion of Vestibular, Optokinetic, Podokinesthetic, and Cognitive Information. Experimental Brain Research. 2006;174(3):528–543. doi:10.1007/s00221-006-0486-7.

73. Pickering SJ. The Development of Visuo-Spatial Working Memory. Memory. 2001;9(4–6):423–432. doi:10.1080/713755973.

74. Goble DJ, Mousigian MA, Brown SH. Compromised Encoding of Proprioceptively Determined Joint Angles in Older Adults: The Role of Working Memory and Attentional Load. Experimental Brain Research. 2012;216(1):35–40. doi:10.1007/s00221-011-2904-8.

75. Quak M, London RE, Talsma D. A Multisensory Perspective of Working Memory. Frontiers in Human Neuroscience. 2015;9:197. doi:10.3389/fnhum.2015.00197.

76. Morris SL, Foster CJ, Parsons R, Falkmer M, Falkmer T, Rosalie SM. Differences in the Use of Vision and Proprioception for Postural Control in Autism Spectrum Disorder. Neuroscience. 2015;307:273–280. doi:10.1016/j.neuroscience.2015.08.040.

77. Izawa J, Pekny SE, Marko MK, Haswell CC, Shadmehr R, Mostofsky SH. Motor Learning Relies on Integrated Sensory Inputs in ADHD, but Over-selectively on Proprioception in Autism Spectrum Conditions. Autism Research. 2012;5(2):124–136. doi:10.1002/aur.1222.

78. Haswell CC, Izawa J, Dowell LR, Mostofsky SH, Shadmehr R. Representation of Internal Models of Action in the Autistic Brain. Nature Neuroscience. 2009;12(8):970–972. doi:10.1038/nn.2356.

